# Extended maturation of the HD10.6 immortalised human dorsal root ganglion cell line enables modelling of nociceptive responses and neural injury

**DOI:** 10.64898/2026.06.01.729276

**Authors:** Kiran Dhillon, Mosab Ali Awadelkareem, Jimena Perez-Sanchez, Geogios Baskozos, David LH Bennett

## Abstract

Human sensory neuron models are an important resource for studying pain mechanisms and axon injury and repair. Current systems are limited by accessibility, scalability, or incomplete functional maturation. The HD10.6 human dorsal root ganglion-derived immortalised cell line represents a promising alternative; however, its maturation trajectory and suitability for disease modelling remain incompletely defined.

Here, we performed a longitudinal, multi-modal characterisation of HD10.6 cells during differentiation over 28 days. Bulk RNA sequencing revealed progressive transcriptional remodelling, with temporal up-regulation of neuronal and nociceptor-associated gene programmes, including ion channels implicated in pain signalling. Protein-level analyses confirmed increased expression of key nociceptor markers and neuropeptides, including TRPV1, Nav1.7, Nav1.8 and CGRP.

Functional assays demonstrated the emergence of sensory neuron-like properties over time. Calcium imaging revealed increasing responsiveness to capsaicin, allyl isothiocyanate, α,β-MeATP, and prostaglandin E2, while patch-clamp electrophysiology at DIV 21 after maturation showed repetitive firing of action potential and, most importantly, exhibited TTX-Resistant sodium currents. These findings establish a temporal relationship between transcriptional changes and functional competence.

Finally, we evaluated the utility of HD10.6 neurons for modelling axon degeneration. Treatment with vacor-induced robust neurite degeneration, which was attenuated by pharmacological inhibition of SARM1, demonstrating engagement of conserved axon degeneration pathways.

Together, our findings define the progressive maturation of HD10.6 sensory neurons and establish this system as a scalable human platform for studying nociceptor biology and SARM1-dependent axon degeneration.

## 1. Introduction

Chronic pain affects at least one in five of the population, and prevalence is predicted to increase with the aging population, diabetes epidemic and increased cancer survival (Breivik et al., 2006; Baskozos et al., 2023). To understand the mechanisms underlying chronic pain and test new molecular treatment targets, we need better human cellular models of pain (Middleton et al., 2021). Nociceptors are sensory neurons that detect injurious stimuli and studying their function and modulation in vitro has been a key component of pain research. This has occurred in concert with our improved understanding of the molecular identities of the transducers and transmission of noxious stimuli (Bennett et al., 2019; Dubin & Patapoutian, 2010). Although there is a broad similarity in the subtypes of sensory neurons between humans and rodents, there are also important differences in terms of expression profiles; at the level of individual ion channels, there are significant cross-species differences in function (Han et al., 2015). Human cellular models can be a helpful step in translation, for instance, in testing treatment effects, including a personalised approach (Lampert et al., 2020; Cao et al., 2016) and optimising biomarkers that may be used in clinical trials (Clark et al., 2021).

Over the past 15 years, the use of human induced pluripotent stem cell (iPSC)–derived nociceptors in pain research has expanded substantially, providing a scalable and human-relevant source of sensory neurons. Early differentiation protocols (Chambers et al., 2012) enabled the generation of sensory neuron–like cells, while subsequent refinements (Deng et al., 2023) improved neuronal maturity and subtype specification. Further advances incorporating co-culture systems with Schwann cells and satellite glial cells (Clark et al., 2017; LeBlang et al., 2025) have enhanced the modelling of myelination and axon–glia interactions. More recently, the development of dorsal root ganglion (DRG) organoids and assembloids (Kim et al., 2025; Mazzara et al., 2020) has enabled more physiologically relevant in vitro models of human pain.

These models have provided important information on the functional properties of human nociceptors and disease pathogenesis. Challenges remain in that these iPSC -derived nociceptors do not model the full range of sensory neuron sub-populations, some channels which have a key role in nociceptors (such as the TTX-resistant voltage-gated sodium channel NaV1.8 (Eberhardt et al., 2015) are not well represented in certain differentiation protocols, and when using small molecules to regulate a developmental trajectory, there is often variation between differentiations in addition to donor-to-donor variation (Schwartzentruber et al., 2018). There is greater availability of native human DRG neurons removed, for instance, during spinal surgery (P. R. Ray et al., 2023) from live donors or from donors post-mortem (Yu et al., 2024). Using native human DRG neurons has the advantage of the full array of sensory neuron subpopulations and the full mature (i.e. adult) molecular profile of nociceptors. However, the scarcity of this resource remains a limiting factor and although the impact of common disorders (such as diabetic neuropathy) on the properties of native human DRG neurons these neurons are not available for rare disorders. There are now established pipelines for the development and biobanking of iPSCs from human participants with rare genetic disorders impacting on nociceptor function (such as the voltage-gated sodium channel variants function (Bennett et al., 2019; Lampert et al., 2020; Van Lent et al., 2024)

The HD10.6 cell line is a conditionally immortalised human DRG-derived neuronal model that carries a tetracycline-suppressible *v-Myc* oncogene that was originally developed from first trimester, embryonic human DRG (Raymon et al., 1999), enabling proliferation in an immature, ‘naive’ state and subsequent maturation into a sensory neuron-like phenotype upon tetracycline treatment. These neurons were reported to have numerous molecular features of nociceptors when studied after one week of maturation, including expression of the transcription factor DRG11, the neuropeptide Substance-P and the NGF receptor trkA. These neurons were excitable, express key transduction elements such as TRPV1 and P2XR3 showed responses to the algogens adenosine triphosphate (ATP) and capsaicin (Y.-C. Chen et al., 2026; Raymon et al., 1999). Hitherto, these neurons have been studied predominantly at relatively early stages of maturation (1 week), and there is a question about their maturity, given that TTX-sensitive sodium currents have been reported, but TTX-resistant currents (mediated by the Nav 1.5, 1.8 and 1.9) have not.

In addition to assessment of acute responses to noxious stimuli, given that peripheral neuropathic pain arises from injury to the peripheral nerves, such platforms can also be used to model the impact of traumatic, toxic (for instance chemotherapy), infective, immune or metabolic challenges. These neurons have already been used to model the impact of herpes simplex virus type-1 on sensory neurons (Q. Zhang et al., 2020).

In this study, we provide a comprehensive characterisation of HD10.6 neurons to assess their suitability as a scalable and reproducible platform for high-throughput pharmacological screening in the context of pain and axonal degeneration. We have looked over a 28-day time course of differentiation and assessed their molecular and functional profile where appropriate drawing comparison with other cellular platforms such as human iPSCd-sensory neurons.

## 2. Results

### 2.1 HD10.6 cells acquire a heterogeneous, DRG-like sensory neuronal transcriptional profile after maturation

HD10.6 cells were expanded in proliferation media and exhibited a doubling time of 28.8 hours (**Fig. S4A**), validating the original measurement of 1.2 days (Raymon et al., 1999). Upon addition of 1 µM tetracycline via replacement with maturation medium 24 hours after seeding, we saw expected growth arrest, although some proliferation still occurred after 48 hours, measured at 72 hours post seeding (**Fig. S4B**).

Recent work has determined transcriptomic profile of HD10.6 sensory neurons to DIV7 (Al-Abbasi et al., 2025). To define transcriptional changes accompanying prolonged HD10.6 maturation, we performed bulk RNA-seq at time points up to 28 days for comparison to undifferentiated HD10.6 cells (UHD), from three independent maturations. Maturation of HD10.6 cells drove a progressive acquisition toward a sensory neuronal transcriptional state. Principal component analysis (PCA) demonstrated a clear separation of UHD cells from cultures matured for 7, 14, 21, or 28 days, with these samples distributed along a temporal maturation trajectory on PC1 (**Fig. 1A**). Notably, HD28 samples showed wider variation within group.

**Figure 1:**
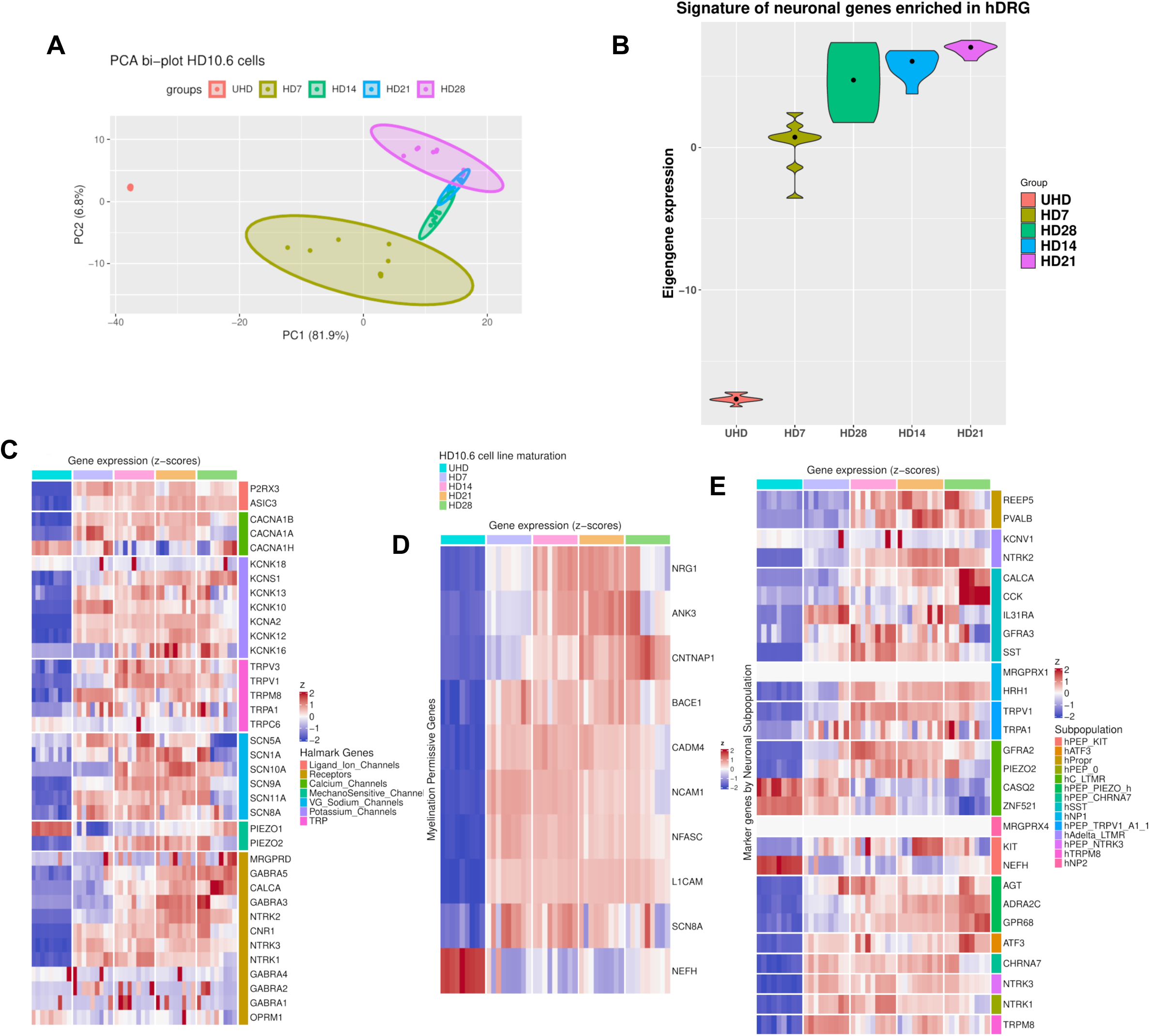
Transcriptomic maturation of HD10.6 cells into sensory neurons. **(A)** Principal component analysis (PCA) of RNA-seq data from immature HD10.6 cells (UHD) and HD10.6 sensory neurons matured for 7 days (HD7), 14 days (HD14), 21 days (HD21) and 28 days (HD28). Differential Expression analysis results and enrichments for biological processes in supplemental figure 1 **(B)** Violin plot showing quantification of eigengene expression of a human dorsal root ganglion (hDRG)-enriched neuronal gene signature (Ray et al., 2018) across UHD, HD7, HD28, HD14 and HD21 samples. **(C)** Heatmap showing expression (z-scores) of selected hallmark sensory neuronal genes (Barry et al., 2023) across the HD10.6 maturation. **(D)** Heatmap showing expression (z-scores) of selected myelination-permissive genes across the HD10.6 maturation time course, including genes implicated in axon–glia interaction and myelination competence. **(E)** Heatmap showing expression (z-scores) of selected neuronal subtype-associated marker genes across the HD10.6 maturation. Genes were grouped according to annotated neuronal subpopulations determined by (Yu et al., 2024). Expression of individual markers in supplemental figure 2. Data are shown for independent RNA-seq samples at each maturation stage (n = 9 per group; 3 technical replicates from 3 independent maturations). In heatmaps, rows represent genes and columns represent samples, with colours indicating relative expression after z-score normalisation.

Consistently, eigengene analysis based on a curated neuronal signature derived from bulk RNA-seq of human DRGs (P. Ray et al., 2018) demonstrated progressive enrichment with HD10.6 maturation, reaching highest levels at HD21, although again, broader distribution at HD28 indicated greater within-group heterogeneity (**Fig. 1B**).

Hallmark sensory neuronal genes (Barry et al., 2023), including *TRPV1*, *TRPA1*, *SCN10A*, *SCN11A*, *PIEZO2* and *NTRK1/2/3*, were progressively upregulated during maturation (**Fig. 1C, E**), together with genes associated with neuronal structure and axon-glial interactions, including *NRG1*, *CNTNAP1* and *NFASC*. One exception to this was that *NEFH* gene (encoding neurofilament heavy chain) which was present in the undifferentiated state and initially down-regulated on differentiation although expression increased with maturation and we could detect the protein product of this gene (DIV 7-28, see section 2.2). **Fig. 1D**) (Bhuiyan et al., 2024; Yu et al., 2024).

Gene ontology enrichment of upregulated genes revealed strong enrichment for neuronal processes including synapse assembly, axon guidance, GPCR signalling, potassium ion transport, sensory perception of pain and nervous system development (**Fig. S1E–H**).

To place HD10.6 sensory neuron maturation in the context of another *in vitro* model, bulk RNA-seq data from induced pluripotent stem cell derived sensory neurons (iPSCdSNs), differentiated for 4 weeks (young) or 6 months (old) from four independent cell lines (AD2, AD3, AH017, NHDF) was used for comparison after batch-correction. PCA analysis between datasets showed convergence of all neuronal groups along PC1 away from their parental immature/undifferentiated state, showing increased transcriptomic effect of maturation/differentiation (**Fig. 2A**).

**Figure 2:**
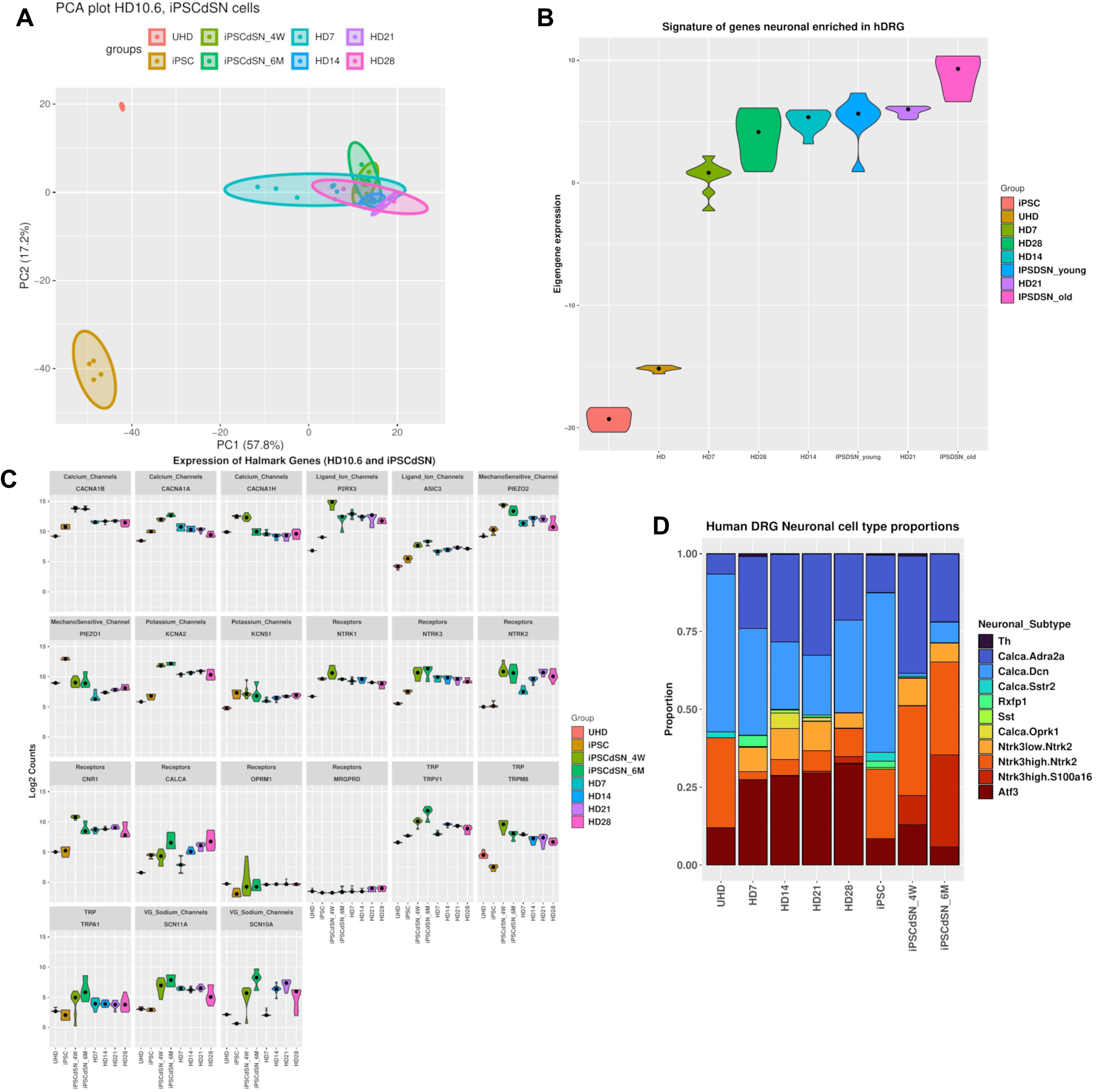
Transcriptomic comparison of matured HD10.6 cells with iPSC-derived sensory neurons identifies shared sensory neuronal features and differences in subtype composition. **(A)** Principal component analysis (PCA) of RNA-seq data from immature HD10.6 cells (UHD), undifferentiated human iPSCs (iPSC), HD10.6 sensory neurons matured for 7 days (HD7), 14 days (HD14), 21 days (HD21) and 28 days (HD28), and human iPSC-derived sensory neurons matured for 4 weeks (iPSCdSN_4W) or 6 months (iPSCdSN_6M) 4 cell lines (AD2, AD3, AH017, NHDF). For comparison, datasets were regularised, log transformed, and batch corrected. **(B)** Violin plot showing eigengene expression of a human dorsal root ganglion (hDRG)-enriched neuronal gene signature (Ray et al., 2018) across iPSC, UHD, HD7, HD28, HD14, iPSCdSN_young (4W), HD21 and iPSCdSN_old (6M) samples. **(C)** Expression of selected hallmark sensory neuronal genes derived from (Barry et al., 2023) across UHD, iPSC, iPSCdSN_4W, iPSCdSN_6M, HD7, HD14, HD21 and HD28. Expression is plotted as log_2_ counts. **(D)** Deconvolution of predicted proportions of bulk-RNAseq HD10.6 and iPSC data to human and mouse dorsal root ganglion (DRG) neuronal subtypes from the single-cell DRG neuronal atlas (Bhuiyan et al., 2024) in HD7, HD14, HD21, HD28, iPSCdSN_4W and iPSCdSN_6M samples. Stacked bar plots show the relative contribution of transcriptionally defined neuronal subtypes. Expression plots of neuronal cell subtype markers and separate deconvolution for human and mouse in supplemental figure 3. Data are shown for independent RNA-seq samples. PCA plots show the first two principal components. In **(C)**, expression values are shown as log_2_ counts. In **(D)**, stacked bars represent relative subtype proportions inferred from cross-species deconvolution.

Additionally, eigengene expression of the hDRG-enriched neuronal signature between iPSC and HD10.6 groups displayed a similar outcome, with iPSCdSN_6M showing the highest level of enrichment, with iPSCdSN_4W sitting within ranges of matured HD10.6 samples (**Fig. 2B**). However, the data suggests that HD10.6 sensory neurons can acquire a mature, enriched transcriptional profile more rapidly than iPSCdSNs. As in **Fig. 1A** and **2A**, HD28 showed broader spread than HD21, indicating that increased late-stage variability is retained in comparative analysis. However, this heterogeneity was also seen in the iPSCdSN samples. The increased expression of sensory neuron hallmark genes was also broadly mirrored in the maturation of both iPSCs and HD10.6 cells with the exception of *MRGPRD* and *OPRM1* which had low levels of expression and showed very little increase in expression with maturation (**Fig. 2C**). Deconvolution of predicted proportions of bulk-RNAseq HD10.6 and iPSC data to human and mouse dorsal root ganglion (DRG) neuronal subtypes from the single-cell DRG neuronal atlas (Bhuiyan et al., 2024) showed that most sensory neuron sub-types were represented (15 out of the 18) HD10.6 D28. Populations which we were unable to detect included: *MRGPRD* and *MRGPRA3* positive C-fiber High-Threshold Mechanoreceptors, cold sensitive *TRPM8* positive C-fibers, *PVALB* proprioceptors and a subset of *CALCA+* Αδ HTMRs *and CHRN7* and *KIT* positive. The composition was similar to iPSCdSNs although the proportions varied (**Fig. 2D**).

### 2.2 HD10.6 sensory neurons express key nociceptive markers

The original developers of the HD10.6 cell line, Raymon et al., reported that these cells exhibited phenotypic features consistent with small-diameter nociceptors. To substantiate and extend this initial observation, we examined the expression of key sensory neuronal markers and nociceptor-associated ion channels across HD10.6 maturation using immunocytochemistry and western blotting (Dykes et al., 2010; Middleton et al., 2021).

Western blotting performed across increasing maturation stages demonstrated progressive acquisition of sensory neuronal proteins (**Fig. 3A-B**). HD10.6 sensory neurons displayed robust expression of BRN3a, a transcription factor enriched in sensory neurons, from DIV (days in vitro) 7 onwards, with expression maintained across later maturation stages and broadly comparable to that observed in iPSCdSNs, which we differentiated using the ‘Chambers’ differentiation protocol (Chambers et al., 2012)(**Fig. 3A, C**). In contrast, immature HD10.6 cells lacked detectable BRN3a expression, consistent with their proliferative, non-neuronal phenotype. Expression of the nociceptor-associated receptor TRPV1 was similarly low in immature cells, lacking the lower kDa band at approximately 97, but increased markedly with maturation, particularly from DIV14 onwards, and remained elevated through DIV21-DIV28 (**Fig. 3 A, C**).

**Figure 3:**
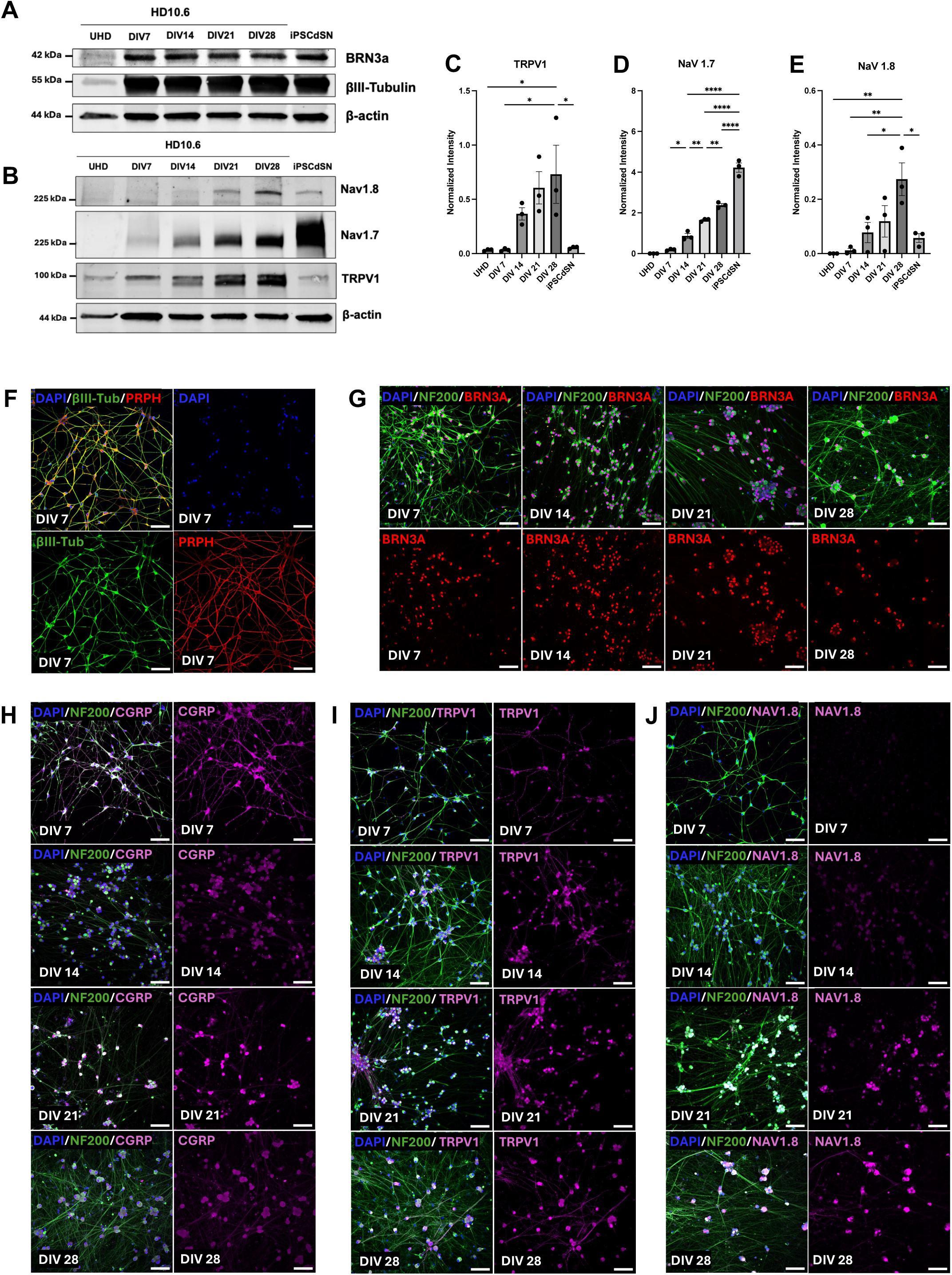
Protein expression of HD10.6 sensory neurons across maturation. **(A - B)** Representative western blots showing expression of neuronal (bIII-tubulin) and sensory neuron-associated markers (BRN3a, TRPV1, NaV 1.7 and NaV 1.8) of immature HD10.6 (UHD) and matured HD10.6 sensory neurons at Day in Vitro (DIV) 7, DIV 14, DIV 21 and DIV 28, and iPSC-derived Sensory neurons of DIV 50-55, from three separate biological replicates. **(C - E)** Quantification of TRPV1, NaV1.7 and NaV1.8 western blot data shown in **(A)**, where intensity is normalised to the corresponding loading control. Bars represent mean ± SEM (N = 3) Ordinary one-way ANOVA with Tukey’s post hoc. **(F)** Representative immunofluorescence images of HD10.6 sensory neurons at DIV 7 stained for βIII-tubulin (green) and peripherin/PRPH (red), with DAPI (blue) nuclear counterstain, shown as merged image. **(G)** Representative immunofluorescence images of HD10.6 sensory neurons stained for NF200 (green) and BRN3A (red), with DAPI (blue), at DIV 7, DIV 14, DIV 21 and DIV 28. Corresponding BRN3A single-channel images are shown below each merged panel. **(H)** Representative immunofluorescence images of HD10.6 sensory neurons stained for NF200 (green) and CGRP (magenta), with DAPI (blue), at DIV 7, DIV 14, DIV 21 and DIV 28. Corresponding CGRP single-channel images are shown for each time point. **(I)** Representative immunofluorescence images of HD10.6 sensory neurons stained for NF200 (green) and TRPV1 (magenta), with DAPI (blue), at DIV 7, DIV 14, DIV 21 and DIV 28. Corresponding TRPV1 single-channel images are shown for each time point. **(J)** Representative immunofluorescence images of HD10.6 sensory neurons stained for NF200 (green) and NaV1.8 (magenta), with DAPI (blue), at DIV 7, DIV 14, DIV 21 and DIV 28. Corresponding NaV1.8 single-channel images are shown for each time point. Scale bar = 100 μm.

We additionally assessed the expression of the nociceptor-associated voltage-gated sodium channels NaV1.7 (encoded by SCN9a) and NaV1.8 (encoded by SCN10a). Western blotting showed progressive upregulation of NaV1.7 from DIV7 to DIV28, with highest levels observed at DIV28, although expression remained greater in DIV54 iPSCdSNs (**Fig. 3B, D**). NaV1.8 was also detected in matured HD10.6 cultures, with signal increasing across maturation and peaking at DIV28, although at lower overall abundance and with greater variability than NaV1.7 (**Fig. 3B, E**). There was some heterogeneity in terms of the level of expression between replicates however the increased expression of TRPV1, NaV1.7 and NaV1.8 broadly mirror the changes at transcript level shown above and at TRPV1 and NaV1.8 could be more reliably detected in DIV28 HD10.6 cells than in DIV54 iPSCdSNs. Together, these data indicate that HD10.6 cells progressively acquire expression of proteins associated with sensory neuronal identity and nociceptor function.

Immunofluorescence staining further confirmed sensory neuronal maturation of HD10.6 cultures. At DIV7, HD10.6 sensory neurons exhibited extensive βIII-tubulin-positive neurite networks and clear expression of peripherin (PRPH), consistent with a peripheral sensory neuronal phenotype at an early stage of differentiation (**Fig. 3F**). Across DIV7-DIV28, cultures showed widespread co-expression of NF200 and BRN3a, and displayed the expected subcellular distribution and nuclear localisation of BRN3a (Dykes et al., 2010), confirming sustained neuronal differentiation and sensory lineage specification throughout maturation (**Fig. 3G**),

Expression of nociceptor-associated neuropeptides and ion channels was also evident. Substance P immunoreactivity has already previously been detected (Dochnal et al., 2026). CGRP expression was present in neuronal cell bodies and processes across the maturation series (**Fig. 3H**). TRPV1 staining was also clearly detectable and became more pronounced with maturation (**Fig. 3I**), consistent with the western blot data. To further elucidate nociceptor-associated sodium channel expression, DIV7-DIV21 HD10.6 sensory neurons were stained for the TTX-resistant sodium channel NaV1.8, mutations of which have been implicated in painful neuropathies (Faber et al., 2012) and showed positive immunoreactivity that was more evident at later stages (**Fig. 3J**).

Notably, morphological changes were also evident across maturation. While early cultures already exhibited extensive neurite outgrowth, later-stage HD10.6 sensory neurons showed more clustered cell bodies, with a tendency towards smaller, rounder somata and increased neurite density, particularly when comparing NF200 staining at later timepoints to DIV7 (**Fig. 3G-J**).

Collectively, these findings show that HD10.6 cells acquire hallmark molecular and morphological features of maturing nociceptor-like sensory neurons, including expression of sensory lineage markers, neuropeptides and nociceptor-associated ion channels.

### 2.3 Electrophysiological properties of matured HD10.6 sensory neurons

The electrophysiological properties of HD10.6 sensory neurons were obtained by whole cell-patch clamp recordings at DIV21, which displayed a stable, mature phenotype from characterisation thus far.

Table in **Fig. 4G** summarises the passive membrane properties measured. We found an average cell capacitance of 16.0 ± 0.8 pF, consistent with the small size of these cells (Dochnal et al., 2026). In the absence of applied current, these cells exhibited stable RMP between -45 mV and -65 mV, as previously reported (Dochnal et al., 2026). HD10.6 exhibited high R_in_ (595.7 ± 21.2 MΩ) and readily fired action potentials when short depolarising pulses were applied (**Fig. 4A**; Rheobase: 34.5 ± 2.6 pA). We also found that many cells fired multiple action potentials in response to prolonged depolarisations (**Fig. 4B**). However, most cells did not maintain repeated firing at very large depolarisations (**Fig. 4C**).

**Figure 4:**
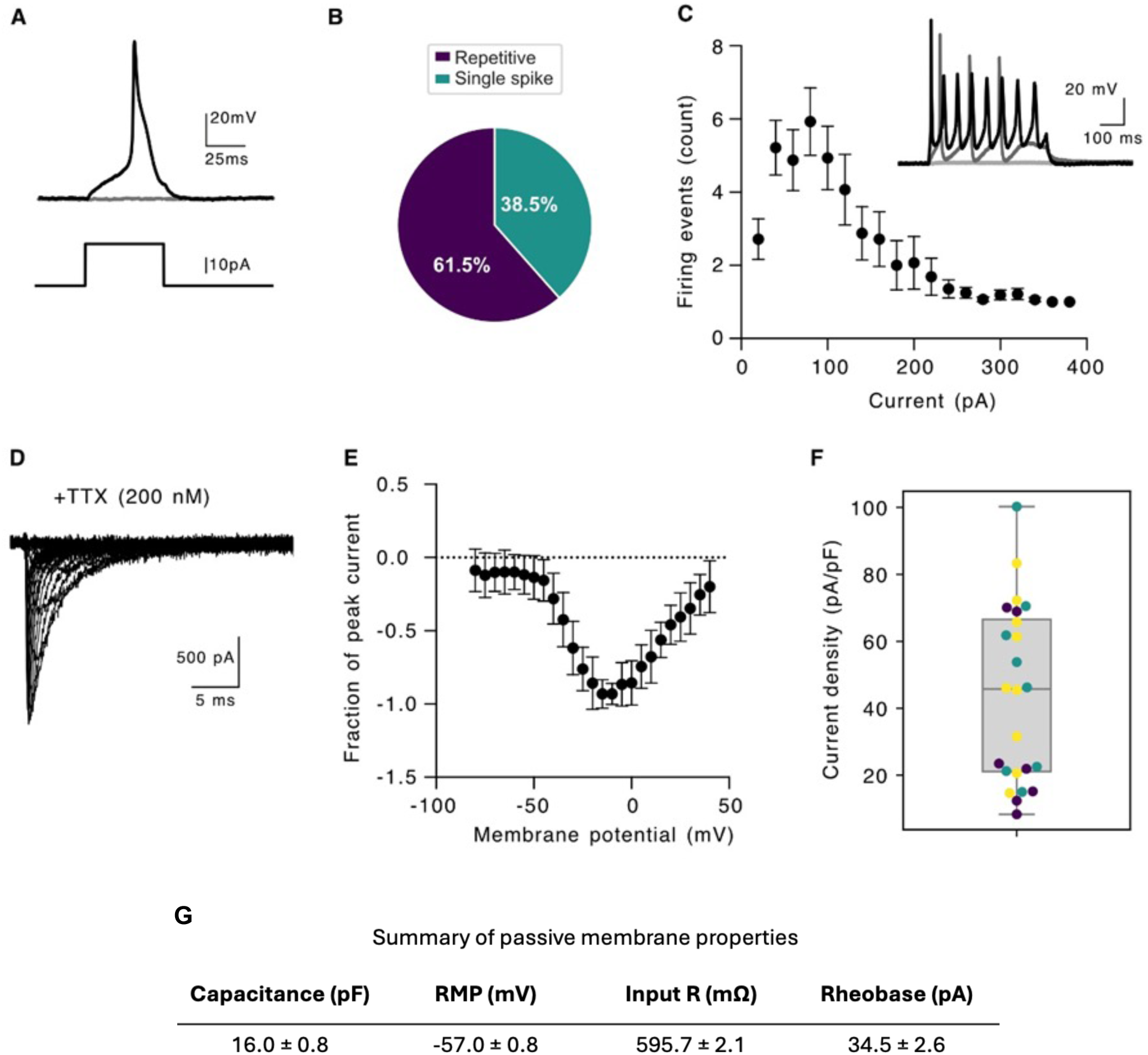
Electrophysiological properties of HD10.6 cells after 21 days of maturation. **(A)** Representative trace showing an action potential elicited in DIV 21 HD10.6 sensory neurons at rheobase (20 pA). **(B)** Percentage of HD10.6 neurons firing single vs. multiple action potentials upon prolonged depolarising current injections. **(C)** Summary of supra-threshold firing in HD10.6 sensory neurons showing repetitive firing. Note that most cells will fire a single action potential with large current injections. Inset shows representative traces of repetitive firing with current steps of 20 pA (grey) and 100 pA (black). **(D)** Representative trace of voltage-gated sodium current induced by step depolarization from a holding potential of −100 mV in the presence of 200 nM TTX. **(E)** Pooled current–voltage (I-V) relationships from HD10.6 sensory neurons (n = 24). **(F)** Quantification of current densities at maximum peak current (n = 24). **(G)** Table showing Membrane properties of HD10.6 cells obtained by whole-cell patch-clamp. (n= 26)

While many voltage-gated sodium channel (VGSCs) isoforms are expressed in sensory neurons, native nociceptors are characterised by the expression of VGSCs resistant to tetrodotoxin (TTX-R). We measured these currents in HD10.6 cells in the presence of 200 nM TTX and found that many cells exhibited TTX-R currents, with a peak current amplitude around -10 mV (**Fig. 4D-E**). Peak densities measured at the maximum current averaged 43.85 ± 5.4 pA/pF but included cells with negligible currents (**Fig. 4F**).

Overall, we found that HD10.6 cells are excitable at DIV21 after maturation, showing repetitive firing. Importantly, these cells express TTX-R currents. This suggests that these cells express NaV1.8, consistent with their protein expression (**Fig. 3B and J**), although we cannot exclude the possibility that that these currents are mediated by other TTX-R channels.

### 2.4 HD10.6 sensory neurons respond to nociceptor-attributed sensory stimulants, increasing with maturation age

To investigate and characterise functionality of any maturation-dependant shifts in HD10.6 sensory neurons, we evoked and measured calcium (Ca^2+^) response using various sensory stimulants at DIV7, 14, 21 and 28 (**Fig. 5**). We used depolarisation in response to potassium chloride (KCl) as a benchmark of excitable cells and the amplitude of the ΔF/F₀ Ca^2+^ response increased with maturity from day 7 to 28.

**Figure 5:**
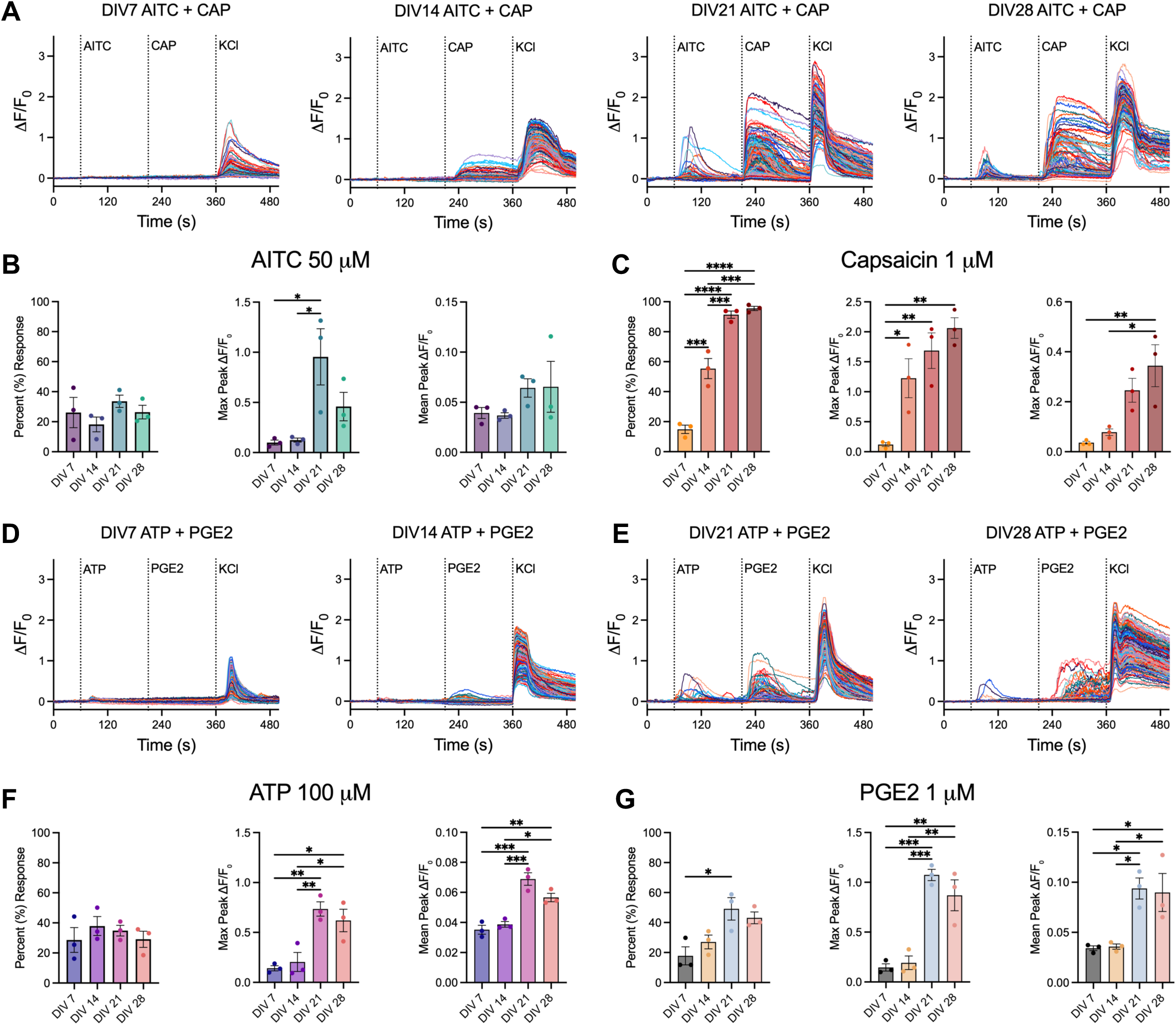
Functional assessment of HD10.6 sensory neurons with calcium imaging through maturation. **(A)** Representative calcium responses (ΔF/F₀) of matured HD10.6 neurons at DIV7, DIV14, DIV21 and DIV28 following sequential application of 50 µM allyl isothiocyanate (AITC), 1 µM capsaicin (CAP) and 50 mM KCl, with timings indicated by vertical dashed lines. **(B)** Quantification of AITC-evoked responses across maturation, shown as percentage of responding cells, maximal peak ΔF/F₀, and mean peak ΔF/F₀. (n = 2453 (DIV7), 2239 (DIV14), 2595 (DIV21), 1445 (DIV28) cells from 3 independent maturations) **(C)** Quantification of capsaicin-evoked responses across maturation, shown as percentage of responding cells maximal peak ΔF/F₀, and mean peak ΔF/F₀. (n = 2157 (DIV7), 1895 (DIV14), 2228 (DIV21), 1722 (DIV28) cells from 3 independent maturations) **(D)** Representative calcium responses at DIV7, DIV14, DIV21 and DIV28 following sequential application of 100 µM α,β-MeATP, 1 µM prostaglandin E2 (PGE2) and 50 mM KCl. **(E)** Quantification of α,β-MeATP-evoked responses across maturation, shown as percentage of responding cells, maximal peak ΔF/F₀, and mean peak ΔF/F₀. (n = 2157 (DIV7), 1895 (DIV14), 2228 (DIV21), 1722 (DIV28) cells from 3 independent maturations) **(F)** Quantification of PGE2-evoked responses across maturation, shown as percentage of responding cells, maximal peak ΔF/F₀, and mean peak ΔF/F₀. (n = 2157 (DIV7), 1895 (DIV14), 2228 (DIV21), 1722 (DIV28) cells from 3 independent maturations) Responding cells threshold: > 3 x SD of pre-30 seconds baseline. For quantification graphs, bars show mean ± SEM and dots represent individual biological replicates/independent experiments. Statistical comparisons are indicated on the graphs; *P < 0.05, **P < 0.01, ***P < 0.001, **\*\*\*\***P < 0.0001.

Cells treated with 1 µM capsaicin displayed increasing response as maturation age progressed (**Fig. 5A**), with a majority of cells responding by DIV21 (**Fig. 5B**), and max ΔF/F₀ mean peak amplitude obtained by DIV28. HD10.6 was originally selected for its TRPV1 sensitivity (Raymon et al., 1999), and as expected we found that nearly all of these cells respond to TRPV1 agonist capsaicin. However, our results indicate that maximal functional response and acquisition of TRPV1 sensitivity was not reached until DIV28.

In contrast, the highest responsive group to treatment with 50 µM allyl isothiocyanate (AITC) was revealed as DIV7 (approximately 66% (Fig. 5B) with this number decreasing to around 44% at DIV14. Response percentage climbed to near DIV7 levels as maturation progressed, where ΔF/F₀ mean peak amplitude was observed at DIV21. The amplitude of AITC responses were measurably much smaller than that for capsaicin.

To understand the changes in response over time, we analysed number of cells responding to AITC only, AITC + Capsaicin and Capsaicin only (**Fig. S5E**), and found ratio between these 3 shifted as cells matured. AITC-only responsive cells were most prevalent at early maturation, whereas acquisition of dual AITC + capsaicin or capsaicin-only sensitivity and responsiveness increased with age. By DIV21-28, the population was dominated by dual ATIC + capsaicin, or capsaicin only responders, with the smallest proportion of AITC-only responding cells.

Our initial investigation (not shown) with 20 µM α,β-Methylene adenosine 5′triphosphate (α,β-MeATP) saw little response at any maturation age of the HD10.6 sensory neuron cultures, however increasing temporal resolution and stimulation dose to 100 µM exhibited a strong response (**Fig. 5E**). The highest response to α,β-MeATP was observed in DIV7 (approximately 70%) and DIV14 (approximately 67%) populations and lessened by DIV28 (**Fig. 5F**). The highest ΔF/F₀ mean peak amplitude was observed in DIV21 neurons (**Fig. 5F**).

Treatment with 1 µM Prostaglandin E2 (PGE2) evoked similar proportion of responses across all maturation timepoints, with slightly higher percentages seen at DIV21 and 28 (**Fig. 5E**). However, the strongest ΔF/F₀ mean peak amplitudes were observed at the later ages, almost double to that of DIV7 and 14, with DIV21 displaying the strongest response (**Fig. 5G**).

KCl-evoked depolarisation exhibited the smallest ΔF/F₀ mean peak amplitudes at DIV7, with markedly stronger activity at DIV21 and DIV28, consistent with increased neuronal excitability during later maturation ages. DIV28 cultures frequently exhibited spontaneous and PGE2 stimulus-evoked burst-like synchronised events (**Fig. 5C**), which were not observed at earlier timepoints.

We investigated further possible polymodal functionality with treatment of DIV28 neurons with 100 µM menthol. We observed approximately 50% of DIV28 neurons showing robust response to menthol treatment (**Fig. S5H**). As menthol and capsaicin were not tested sequentially in the same cells, we cannot directly infer that the same cells are responsive to both.

Recent work (Dochnal et al., 2026) has suggested that absence of Forskolin, a cAMP agonist (Seamon et al., 1981) in the maturation media led to lower resting action potential of HD10.6 cells. To test any affect to sensory-evoked Ca²⁺ responses, we cultured the HD10.6 sensory neurons without Forskolin and applied the same sensory stimulants at DIV28. However, we found no significant differences between groups for either capsaicin and KCl for percentage of cells responding or mean ΔF/F₀ peak amplitude across three independent biological repeats (**Fig. S5C-D**).

Additionally, HD10.6 sensory neurons expressing fluorescent calcium indicator, GCaMP8f display calcium transients which can be captured and quantified as a proxy for spontaneous activity (**Fig. S6A**). DIV7 HD10.6 sensory neurons were transduced with *AAV-CAG-jGCaMP8f-WPRE* (Y. Zhang et al., 2023), kept in culture until DIV32, and live fluorescence was recorded at 10 Hz over 30 seconds. Live dead staining of neurons displayed no co-localisation with GCaMP8f fluorescence, confirming expression present only in live cells (**Fig. S6C**). Spontaneous calcium transients could be observed and treatment with PGE2 (1 µM) resulted in an increase of GCaMP8f activity, when integrated fluorescence between baseline and post-treatment groups is compared (**Fig. S6B**).

Taking together the data obtained across all stimuli tested, we observed the most robust and consistent responses at DIV21-28, suggesting that HD10.6 sensory neurons require at least three weeks of maturation to achieve a stable, functional response to nociceptive sensory calcium stimulants.

### 2.5 Myelination of HD10.6 sensory neurons with Rat Schwann cells

Co-culture in pro-myelination conditions (Clark et al., 2017) of HD10.6 sensory neurons with Rat Schwann cells (RSCs) at maturation day 7, results in myelination (**Fig. 6A**). Immunostaining for Myelin Basic Protein shows myelin formation within 3 weeks of co-culture. RSCs present a pro-myelinating phenotype with confirmation via immunocytochemistry of transcription factor expression, KROX20 which is a key regulator of myelination (Topilko et al., 1994) (**Fig. 6A**).

**Figure 6:**
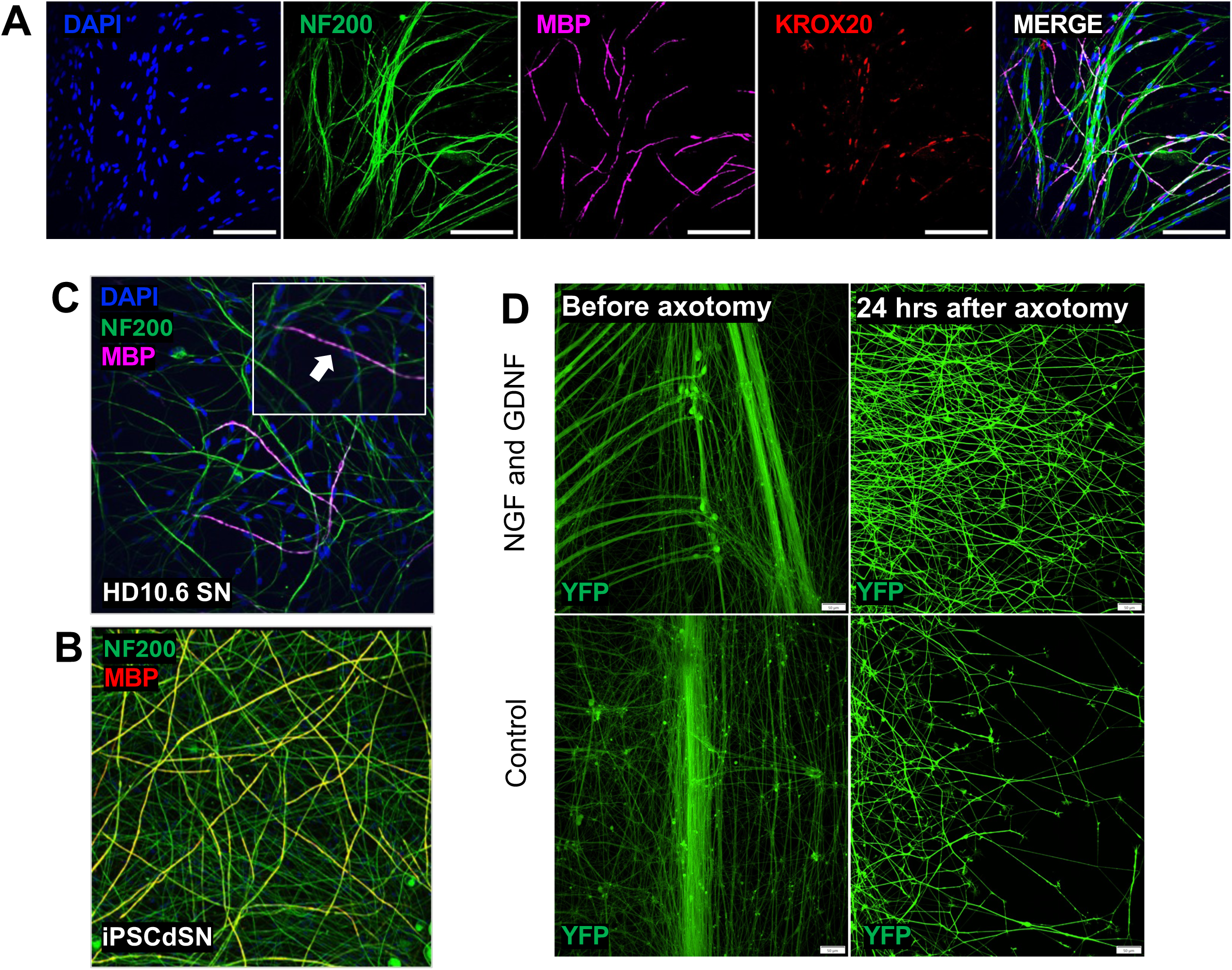
Myelination of matured HD10.6 sensory neurons following co-culture with rat Schwann cells. **(A)** Representative immunofluorescence images of matured HD10.6 sensory neurons following addition of rat Schwann cells at DIV7 and co-culture for 22 days in pro-myelination medium containing tetracycline. Single-channel images for NF200 (green), MBP (magenta) and KROX20 (red), with DAPI nuclear counterstain (blue), and merged are shown. Scale bar = 50 µm. **(B)** Representative immunofluorescence image of myelinated iPSC-derived sensory neurons (iPSCdSN) stained for NF200 (green) and MBP (red), shown as a positive comparison for myelin formation. **(C)** Representative immunofluorescence image of myelinated HD10.6 sensory neurons (HD10.6 SN) stained for NF200 (green), MBP (magenta) and DAPI (blue), showing MBP-positive segments associated with neuronal fibres. Inset shows a higher-magnification view of the boxed region; white arrow indicates irregular myelination. **(D)** Representative live-cell images of EYFP-expressing HD10.6 neurons in the axonal compartment of microfluidic devices before axotomy (pre-injury) and 24 hours post-axotomy under control conditions or following treatment with NGF (50 ng/mL) and GDNF (25 ng/mL). Increased neurite density and extension are observed in NGF/GDNF-treated cultures compared to controls. Scale bar: 50 µm.

However, in contrast to iPSCdSNs which form long, continuous myelin sheaths (**Fig. 6B**) and show positive expression of internodes such as CASPR and Pan-NaV (Clark et al., 2017) myelin formation in HD10.6 is limited to small and sparse segments (**Fig. 6C**).

### 2.6 HD10.6 sensory neurons can be used for studying axonal regeneration

To investigate neurite regenerative capacity in human sensory neurons and to demonstrate that this cell line could be used to screen for pharmacological or genetic interventions that affect regeneration, HD10.6 cells were cultured in microfluidic devices and subjected to axotomy in the axonal compartment at DIV 21, (**Fig. 6D**). Neurite regeneration was assessed 24 hours post-injury by quantifying the area occupied by EYFP-positive neurites and normalising to pre-axotomy values.

Treatment with neurotrophic factors (NGF, 50 ng/mL; GDNF, 25 ng/mL) continuously since DIV 1 resulted in a significant increase in neurite regeneration 24 hours post-injury compared to control conditions lacking growth factors since DIV 1. Representative images demonstrate enhanced neurite density and extension into the axonal compartment in NGF/GDNF-treated cultures relative to controls. Quantitative analysis taken from one maturation experiment (**Fig. S7A)** confirmed a robust increase in the regenerated neurite area in neurotrophic factor treatment culture, indicating that NGF and GDNF promote regenerative outgrowth in HD10.6 human DRG-derived neurons.

### 2.7 HD10.6 sensory neurons can be used for studying Wallerian degeneration

To further demonstrate a higher throughput and scalable use for HD10.6 cells compared to iPSCdSNs or primary rodent DRGs, we developed a model of Wallerian degeneration *in vitro*. A major molecular driver of axon loss is the sterile alpha and TIR motif–containing protein 1 (SARM1). Under axonal stress, depletion of the essential survival factor NMNAT2 leads to accumulation of its substrate nicotinamide mononucleotide (NMN), which allosterically activates SARM1. Once activated, SARM1 rapidly depletes NAD⁺ through its intrinsic NADase activity, precipitating energetic failure, calcium dysregulation, and axonal fragmentation (Essuman et al., 2017; Figley et al., 2021; Gilley et al., 2015). This pathway has been strongly implicated in chemotherapy-induced peripheral neuropathy (CIPN): agents such as paclitaxel, vincristine, and cisplatin converge on a SARM1-dependent degeneration programme, and SARM1 knockout provides robust protection in multiple CIPN models (Bosanac et al., 2021; Geisler et al., 2019). Additional evidence comes from studies using the discontinued rodenticide vacor, whose metabolite vacor-mononucleotide (VMN) potently and specifically activates SARM1, inducing rapid axon degeneration (Loreto et al., 2021).

We first confirmed SARM1 expression in HD10.6 sensory neurons (**Fig. 7A**) via western blot, showing robust expression in the matured groups, in comparison to no expression in the immature state. We subsequently treated cells at DIV28 with increasing doses of vacor, a potent and specific activator of SARM1, to assess cellular health via the CellTitre-Glo assay (Promega), which quantifies metabolically viable cells through ATP-dependant luciferase activity and luminescence intensity measurement. Results showed strong decrease in viability from doses 25 - 100 µM, with little change in viability for lower doses, even after 48 hours treatment (**Fig. 7B**).

**Figure 7:**
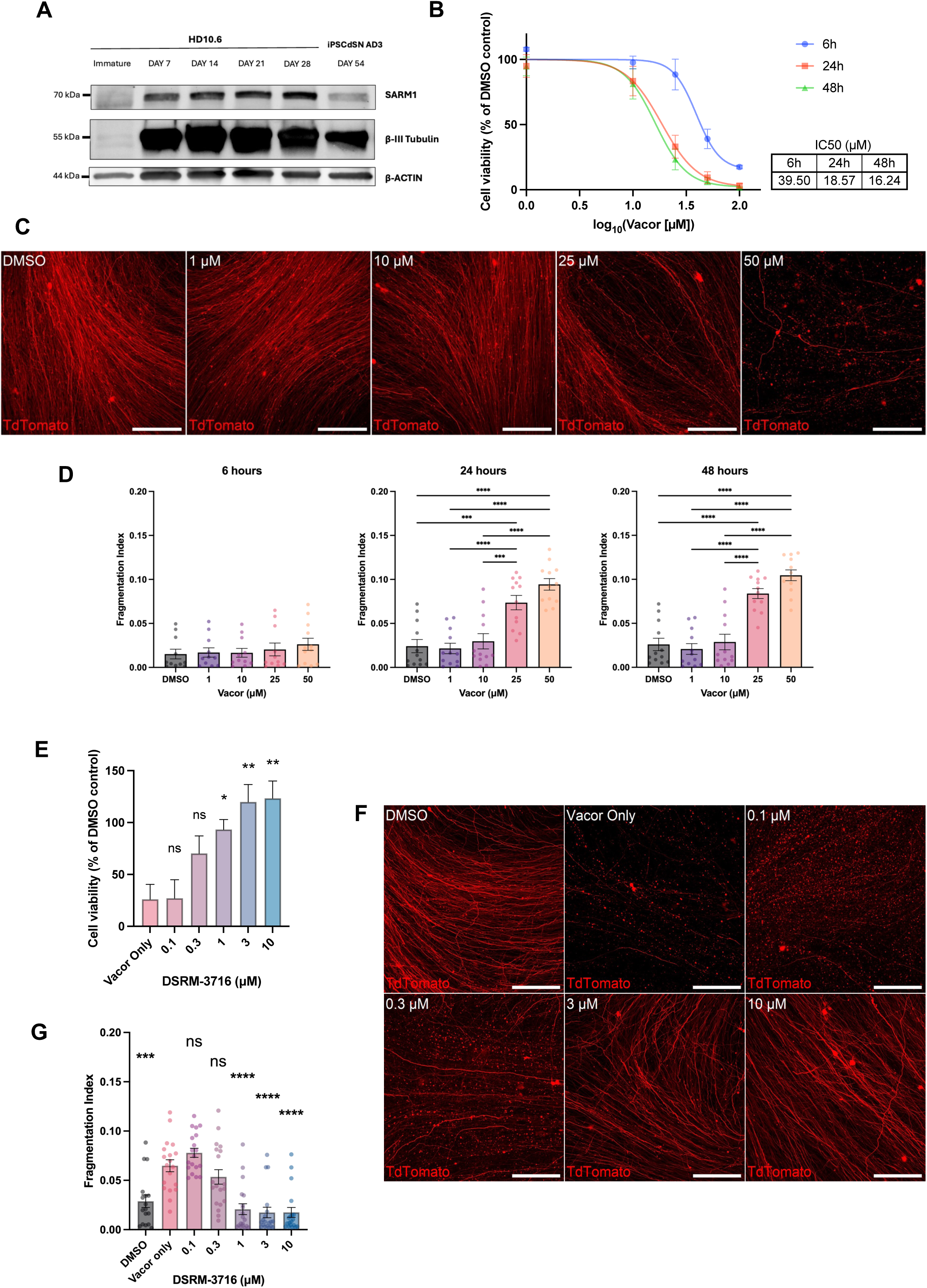
Vacor induced degeneration of matured HD10.6 sensory neurons and rescue by SARM1 inhibition. **(A)** Representative western blot showing SARM1 and βIII-tubulin expression in immature (“naïve”) HD10.6 cells, HD10.6 sensory neurons differentiated to DIV7, DIV14, DIV21 and DIV28, and human iPSC-derived sensory neurons matured for 54 days. **(B)** ATP-based cell viability dose-response curves for DIV28 HD10.6 sensory neurons treated with increasing concentrations of vacor for 6, 24 and 48 h, showing calculated IC50 doses per treatment time. (N = 3) **(C)** Representative live fluorescence images of TdTomato-positive DIV28 HD10.6 sensory neurons treated with increasing concentrations of vacor, showing progressive loss of neurite integrity and fragmentation with increasing dose. **(D)** Quantification of vacor-induced neurite degeneration, expressed as fragmentation index, after 6 h, 24 h and 48 h treatment. **(E)** ATP-based cell viability of DIV28 HD10.6 sensory neurons treated with increasing concentrations of DSRM-3716 in the presence of 25 μM vacor for 24 h. (N = 3) **(F)** Representative live fluorescence images of TdTomato-positive DIV28 HD10.6 sensory neurons treated with DMSO, vacor only, or increasing concentrations of DSRM-3716 in the presence of 25 μM vacor, at 24 h **(G)** Quantification of neurite degeneration in TdTomato-positive HD10.6 sensory neurons co-treated with vacor and DSRM-3716 for 24 h, expressed as fragmentation index Data are shown as mean ± SEM. Statistical analysis was performed using a Kruskal–Wallis test (non-parametric one-way ANOVA) with Dunnet’s post-hoc, with significance as indicated on the graphs; *P < 0.05, **P < 0.01, ***P < 0.001, **\*\*\*\***P < 0.0001; N = 3 independent experiments. Scale bar = 100 μm

To investigate damage and loss of axonal integrity, we next transduced DIV7 HD10.6 sensory neurons with adeno-associated virus, *AAV9-CAG-TdTomato*, to enable visualisation and tracking of vacor-induced fragmentation via live cell imaging.

The TdTomato+ HD10.6 sensory neurons were next co-cultured with Rat Schwann cells at DIV14, after which cultures were sustained in pro-myelination medium until DIV28. Co-cultures were treated with increasing doses of Vacor and DMSO vehicle. Fragmentation was assessed via calculation of a Fragmentation index (**Fig. S8)** which resolves TdTomato positive pixels as ‘Fragmented’ or ‘Unfragmented’ through segmentation after particle size thresholding. Through this live imaging approach, we captured a similar dose response, correlating ATP viability to fragmentation, which complement each other in this model (**Fig. 7C**).

To elucidate rescue of this vacor-induced SARM1 fragmentation, we employed SARM1 inhibitor DSRM-3716 at varying doses (**Fig. 7E**) and co-dosed with base dose of vacor 25 µM (selected as the lowest dose for most significant fragmentation achieved (**Fig. 7D**)) and imaged cultures at 24 hours (**Fig. 7E**). DSRM-3716 co-treatment displayed a rescue of cell viability with increasing doses (**Fig. 7F**), in comparison to the vacor-only treated group.

Fragmentation after treatment lowest dose of DSRM-3716 (0.1 µM) exceeded significantly the fragmentation measured by the base vacor-only concentration (**Fig. 7G**), which coincides with recent findings that this DSRM inhibitor, along with other mechanistically similar base-exchange SARM1 inhibitors show a cytotoxic affect, proposed by their paradoxical activation of SARM1 at these lower doses (Mani et al., 2025). At higher doses however, we observed effective-dose dependent rescue of vacor-induced fragmentation and ATP viability loss, when compared to vehicle only groups.

## 3. Discussion

Human cell–based models of nociceptors are increasingly used to study pain mechanisms, yet each platform presents distinct advantages and limitations. The HD10.6 cell line provides a scalable, easy-to-work-with human DRG–derived system; however, its level of maturation and similarity to native human nociceptors remains incompletely defined. In this study, we provide a comprehensive molecular and functional characterisation of HD10.6 cells across an extended 28-day maturation time course and benchmark some of their molecular properties against human iPSC-derived sensory neurons and data from primary human and mouse DRG tissue.

Previous studies have demonstrated that HD10.6 cells exhibit enrichment of sensory neuron–associated gene expression and functional responsiveness to nociceptive stimuli. For example, Al-Abbasi et al (Al-Abbasi et al., 2025) reported moderate transcriptomic similarity between HD10.6 cells and human DRG nociceptor subtypes, alongside expression of key ion channels and receptors relevant to pain signalling, although canonical nociceptor markers such as *CALCA*, *SCN10A*, and *TRPV1* were expressed at relatively low levels. Similarly, Dochnal *et al*., (Dochnal et al., 2026) showed that HD10.6 neurons respond to algogens and exhibit sensitisation, supporting their functional relevance. However, both studies focused on early maturation stages (7 days (Al-Abbasi et al., 2025) and up to 14 days (Dochnal et al., 2025) which likely represent a more immature neuronal state. We have found that that the molecular and functional response matures with an optimum nociceptor-like phenotype at 21 and 28 days *in vitro*.

By extending maturation to 28 days, our bulk RNA-seq demonstrated progressive maturation from the immature state towards a DRG-like sensory neuronal transcriptional profile, with increasing enrichment of canonical neuronal and sensory gene programmes across maturation including canonical markers of nociceptors. By DIV21 and DIV28, HD10.6 sensory neurons showed strong enrichment of an hDRG-associated neuronal signature (Yu et al., 2024) and broad transcriptomic overlap with iPSCdSNs, indicating that this model achieves a mature sensory neuronal identity within weeks rather than months (Bhuiyan et al., 2024; P. Ray et al., 2018). This comparatively rapid maturation, together with ease of culture and assay scalability, suggests that HD10.6 cells may facilitate more rapid preclinical investigation.

A key finding of the transcriptomic analysis, however, was that later maturation did not converge on a single uniform endpoint. Although DVI28 sensory neurons retained high neuronal signature scores, they also showed greater dispersion in PCA dimensions and broader variability across marker expression and subtype deconvolution analyses than DIV21 sensory neurons. This suggests that prolonged differentiation is accompanied not only by continued linear maturation but also by increased transcriptional heterogeneity. Subtype analyses supported this interpretation, with matured HD10.6 cultures retaining features of multiple sensory neuronal classes. In particular, HD10.6 sensory neurons showed strong representation of peptidergic and nociceptor-like subtypes marked by *CALCA, TRPV1, TRPA1* and *MRGPRX1/4*, together with smaller contributions from mechanosensory or larger-fibre-associated programmes marked by *NTRK2, NTRK3, PIEZO2, KCNV1* and *NEFH* (Bhuiyan et al., 2025; Middleton et al., 2021; Tavares-Ferreira et al., 2022). Importantly, deconvolution using both human and mouse reference atlases yielded broadly concordant patterns, supporting the robustness of this mixed subtype composition (Bhuiyan et al., 2024). These data suggest that HD10.6 cells mature towards a heterogeneous sensory neuronal state rather than a single terminal subtype, with HD28 representing a more compositionally diverse stage than HD21.

The functional characterisation was consistent with the suggested nociceptor-biased identity from the transcriptomic observations. HD10.6 sensory neurons expressed hallmark proteins associated with maturing nociceptors, including CGRP, BRN3a, TRPV1, NaV1.7 and NaV1.8 **(Fig. 3)**, and showed increasing responsiveness to nociceptive agonists over maturation **(Fig. 5)**. By DIV28, capsaicin responsiveness was near-universal, consistent with the original characterisation of HD10.6 as a nociceptor-like human DRG-derived line (Raymon et al., 1999) and with later reviews of its sensory phenotype (Haberberger et al., 2020). This degree of capsaicin responsiveness is lacking in many stem cell-based sensory neuron differentiation protocols, in which TRPV1-positive yields may remain lower or vary between protocols, donors or cell lines (Deng et al., 2023; Fofie et al., 2025; Kalia et al., 2024; Schrenk-Siemens et al., 2015). Early AITC-only responses then shifted towards predominantly dual AITC and capsaicin sensitivity by DIV21-DIV28, consistent with the established overlap between TRPA1- and TRPV1-expressing nociceptors (Middleton et al., 2021; Story et al., 2003). The lower capsaicin response seen at earlier stages despite detectable TRPV1 immunostaining suggests that transcriptional or protein expression may precede full functional channel competence, underscoring the importance of combining phenotypic and functional characterisation. Consistent with TRPM8 expression, a substantial subset of mature HD10.6 responded to menthol acts through TRPM8 receptor (which mediates cool sensing). Menthol-sensitive neurons may overlap with capsaicin-responsive populations *in vitro* (Dhaka et al., 2008; Hjerling-Leffler et al., 2007; Karashima et al., 2007; Moparthi et al., 2022). Electrophysiological recordings demonstrated that mature HD10.6 (at DIV21) exhibit functional excitability, including repetitive firing of action potentials, as has been recently shown (Dochnal et al., 2025). They also display TTX-resistant (TTXR) sodium currents, which are characteristic of nociceptors and are mediated by the VGSCs NaV1.8 and NaV1.9 in mature neurons (Akopian et al., 1996; Cummins et al., 1999). Our recording protocol was not optimised to measure NaV1.9 currents (Dib-Hajj et al., 2002), suggesting that these currents are likely mediated by NaV1.8, although the contribution of NaV1.9 cannot be excluded. Together, these observations indicate that HD10.6 cells acquire a functionally mature nociceptor-like phenotype consistent with the findings on transcriptional profiling.

Comparison with iPSCdSNs suggests the two systems share broad sensory neuronal features but differ in both practical utility and subtype balance. Transcriptomically, HD10.6 sensory neurons appeared relatively enriched for *CALCA*-associated peptidergic nociceptor-like and TRPM8-related sensory populations, whereas iPSCdSNs showed relatively greater representation of NTRK2/NTRK3/PVALB-associated mechanosensory-like populations (Bhuiyan et al., 2024; Middleton et al., 2021). This difference is likely to be biologically meaningful, as peptidergic and nociceptor-like populations are more often associated with smaller-diameter, weakly myelinated or unmyelinated axons, whereas larger-fibre mechanosensory populations are typically more permissive to Schwann cell myelination (Middleton et al., 2021).

This distinction is particularly relevant to the myelination phenotype. The limited myelination observed in HD10.6 co-cultures, especially compared to myelination of iPSCdSNs under similar conditions (Clark et al., 2017; Topilko et al., 1994) is likely to reflect incomplete and heterogeneous acquisition of a myelination-competent neuronal state. Although rat Schwann cells adopted a KROX20-positive pro-myelinating (Ghislain & Charnay, 2006) phenotype (**Fig. 6A**) and HD10.6 neurons expressed key axon–glial and myelination-associated genes, including *NRG1, CNTNAP1, and NFASC.* These programmes arose within a transcriptionally mixed population enriched for nociceptor-like and peptidergic sensory identities. Given the established role of axonal NRG1 in Schwann cell myelination (Brinkmann et al., 2008; S. Chen et al., 2006; Liang et al., 2012), and the importance of axoglial proteins such as CNTNAP1 and NFASC in peripheral nerve organisation, these data suggest that the molecular machinery required for Schwann cell engagement is present but may not be deployed uniformly across the culture. Therefore, it could be plausible that initiating myelination at later stages of HD10.6 maturation may improve efficiency, particularly as myelination-associated genes were more prominent at later timepoints (**Fig. 1D**), although this will require direct experimental testing.

Further to this, HD10.6 sensory neurons presented a higher proportion of *ATF3* expression in comparisons to the iPSCdSNs. Of course, it is likely that stress from increasingly denser cultures through maturation of the HD101.6 sensory neurons did contribute to this *ATF3* upregulation especially in DIV28, and it is plausible that the increased neuronal activity seen at these later time points elevated their metabolic demands, which may warrant more media replenishment than the current protocol used. However, there are interestingly, recent suggestions (Bhuiyan et al., 2025) of an ATF3 phenotype existing as a transitory regenerative state in DRG populations, which raises a possibility that this may be enriched in the HD10.6 sensory neuronal populations.

We demonstrated HD10.6 cells are well suited to degeneration-based assays and higher-throughput set-ups. The cells can be readily cultured in 96-well plates, with more precision in seeding densities due to a predictable growth/doubling rate (**Fig. S4**) and efficiently transduced with adeno-associated viruses such as GCaMP8f (Y. Zhang et al., 2023) (**Fig. S6**) and TdTomato (**Fig. 7**) as well as lentiviral construct pLenti-Synapsin-EYFP-WPRE (**Fig. 6D**), with robust expression. This is particularly advantageous in dense HD10.6 axonal networks, where NF200-based quantification can be challenging, especially if smaller diameter axons are present, which may express less NF200 (Ren et al., 2018; Ruscheweyh et al., 2007), which was consistent to the downregulation of *NEFH* we saw upon maturation **(Fig. 1D)** and observed in our immunofluorescence staining of later-matured HD10.6 sensory neurons (**Fig. 3F-J**). Fluorescent axonal labelling, therefore, provides a tractable platform for automated analysis of axon fragmentation across large image datasets. In this context, co-culture with rat Schwann cells remained valuable even where compact myelin formation was limited, as Schwann cells provided structural support that reduced detachment and enabled more reliable analysis of degeneration. These features support the use of HD10.6 cells in scalable assays of Wallerian degeneration, chemotherapeutic neurotoxicity and pharmacological rescue, complementing iPSCdSNs which remain an established platform for chemotherapy-induced peripheral neuropathy modelling (Schinke et al., 2021; Xiong et al., 2021; Kalia et al., 2024).

We also demonstrate that neurotrophic factor supplementation enhances neurite regeneration in HD10.6 human DRG-derived neurons following axonal injury. Specifically, combined treatment with NGF and GDNF increased neurite regrowth within 24 hours post-axotomy compared to untreated controls. These findings are consistent with the well-established roles of NGF and GDNF in promoting neuronal survival, axonal growth, and regeneration in sensory neurons (Diaz et al., 2026).

The use of a microfluidic platform enabled spatial separation of somatic and axonal compartments, allowing precise induction of axotomy and quantification of regenerative responses. The observed increase in EYFP-positive neurite area suggests that HD10.6 cells retain responsiveness to neurotrophic cues and can serve as a relevant human model for studying axonal regeneration. Future work comparing neurotrophic factor receptor agonists or disease-specific conditions could further validate the utility of this model in translational research.

Several limitations should be considered. As an immortalised cell line, HD10.6 carries an integrated viral oncogene, and the downstream consequences of immortalisation, including altered growth control or other dysregulated cellular behaviours, are not fully defined. More broadly, continuous cell lines can undergo genetic and transcriptional drift over time, which may affect reproducibility if not carefully controlled (Ben-David et al., 2018) a concern typically mitigated in iPSC systems through rigorous quality-control measures such as routine karyotyping (Lund et al., 2012; Steichen et al., 2019; Park et al., 2024). In addition, despite their origin from human embryonic DRG tissue, HD10.6 cells are derived from a single selected clone and therefore cannot capture the full transcriptional, developmental and functional heterogeneity of native human DRG populations *in vivo* (Bhuiyan et al., 2025). Nor can they substitute for patient-derived iPSC models in the study of genotype-specific neuropathies. These limitations should frame the HD10.6 cell line as a fit-for-purpose platform with strengths in reproducibility, accessibility and assay scalability, and a tool to enhance preclinical investigations to complement other models.

Overall, our findings support HD10.6 cells as a robust human nociceptor-oriented sensory neuron model, particularly around DIV21, when the cultures display strong neuronal maturation with less apparent heterogeneity than at later stages. In this window, HD10.6 cells combine transcriptomic, molecular and functional maturity with practical advantages that make them well suited to pharmacological studies, mechanistic investigations and higher-throughput screening in peripheral neuropathy research.

## 4. Methodology

### Cell Culture and maturation of HD10.6 sensory neurons

HD10.6 cells were thawed and seeded on to cultrex or fibronectin coated plates and maintained in proliferation media: Advanced DMEM/F12 (Gibco) supplemented with B-27 without Vit A (Gibco), GlutaMAX (Gibco), Prostaglandin E1 (10 ng/mL), G814 solution (50 µg/mL) and beta-FGF (0.5 ng/mL) at 37 °C with 5% CO₂. To induce maturation into sensory neuronal phenotype, HD10.6 cells were seeded on to poly-d-lysine or poly-l-ornithine and cultrex coated coverslips or plates and cultured in maturation media: Neurobasal medium supplemented with B-27, GlutaMAX, NGF (50 ng/mL), NT-3 (25 ng/mL), GDNF (25 ng/mL), CNTF (25 ng/mL), Forskolin from Coleus (25 µM) and tetracycline (1 µg/mL) at 37 °C with 5% CO₂. Half media changes occurred twice a week. **Table S2** for full reagent details.

### Differentiation of Human iPSCs into Sensory Neurons

Human iPSC lines were obtained through the IMI/EU-sponsored StemBANCC consortium via the Human Biomaterials Resource Centre at the University of Birmingham, UK. The lines were differentiated into sensory neurons using a modified version of the Chambers protocol (Chambers et al., 2012), as previously reported. Briefly, iPSCs were maintained on Matrigel-coated plates in mTeSR1 or StemFlex medium and passaged at around 80% confluency using EDTA. Prior to differentiation, all lines underwent quality control, including assessment of pluripotency markers, genomic integrity, Sendai virus clearance, and mycoplasma testing.

For differentiation, iPSCs were dissociated using EDTA and plated at high density. Neural induction was initiated by culturing cells in knockout serum replacement (KSR) medium (Knockout-DMEM supplemented with 15% knockout serum replacement, 1% GlutaMAX, 1% non-essential amino acids, and β-mercaptoethanol) containing dual SMAD inhibitors SB431542 (10 μM) and LDN-193189 (100 nM). On day 3, additional small molecules were added, including CHIR99021 (3 μM), SU5402 (10 μM), and DAPT (10 μM). Dual SMAD inhibition was withdrawn on day 5, and the medium was gradually transitioned to N2/B27 medium (Neurobasal supplemented with 2% B27, 1% N2, and 1% GlutaMAX) in 25% increments.

At day 11, cells were replated onto coated coverslips or 6-well plate in N2/B27 medium supplemented with BDNF, NGF, NT-3, and GDNF (all at 25 ng/mL). CHIR99021 was maintained for an additional 4 days post-replating. Medium was replaced twice weekly. Where necessary, cytosine β-D-arabinofuranoside (AraC; 1 μM) was added shortly after replating to eliminate proliferating non-neuronal cells and subsequently withdrawn once cultures reached neuronal purity, typically within 2–3 weeks. Cells were maintained with twice-weekly media changes and used for downstream experiments following maturation.

### Growth curves and doubling time

Cell proliferation was quantified using the CellTiter-Glo® Luminescent Cell Viability Assay (Promega). Cells were seeded into 96-well plates at 2,500-15,000 cells per well. As higher densities showed early plateauing of growth within the constraints of the 96-well format, luminescent values for only 2,500-10,000 cells were considered for doubling time calculation. Each seeding density contained a minimum of five technical replicates per time point. After 24 hours of attachment, 1 µM tetracycline was added with full exchange to maturation medium, and cell growth was subsequently assessed at 48 h and 72 h. Doubling time (*Td*) was calculated between the 24 h and 48 h interval, as this period represented the linear/exponential phase of growth based on luminescence curves. The following standard exponential-growth formula was used:

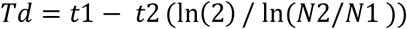

where *t*1 = 24 h and *t*2 = 48 h, and *N*1 and *N*2 represent the mean luminescence values at 24 h and 48 h, respectively, for each seeding density.

### Immunocytochemistry

Cells were fixed in 1% paraformaldehyde (20 min) at room temperature (RT), permeabilised with 0.1% Triton X-100 or ice-cold methanol (for myelinated cultures) and blocked with 5% normal goat serum. Primary antibodies were incubated overnight at 4°C in 0.1% Triton X-100 and 2% normal goat serum in PBS, followed by Alexa Fluor-conjugated secondaries (1:2000, 2 h, RT) in and 2% normal goat serum in PBS. Nuclei were counterstained with DAPI (1 µg/mL).

### Western Blot

HD10.6 cells were lysed in ice-cold RIPA buffer supplemented with protease and phosphatase inhibitors. Protein lysate concentrations were determined using BCA assay. Equal amounts (20 µg per sample) were heated to 95 °C for 5 min, or 75 °C for 15 min for membrane bound proteins, denatured using NuPage reducing agent (Invitrogen) and prepared in Lamelli buffer (Biorad). Samples were resolved on 4 - 20% SDS–polyacrylamide gels (Biorad) via electrophoresis at 80 – 100 mV, and gels transferred to 0.22 µm nitrocellulose membranes using semi-dry TurboBlot system (Biorad). Membranes were probed with primary antibodies overnight at 4°C. Alexa Fluor-conjugated secondary antibodies were incubated for 2 h at RT (1:2000) with Tris Buffered Saline with 0.1% Tween-20 used for wash steps. Fluorescence detection was achieved with the ChemiDoc Imaging System (Biorad). Images were analysed using ImageLab software (Biorad). Fluorescent band intensity was measured with background subtraction, and target proteins were normalised with relative loading controls to determine normalised expression.

### Bulk mRNA-seq

Total RNA was isolated of HD10.6 cells using the RNeasy Mini kit (Qiagen), following protocol instructions including on-column, DNase I treatment and resolved in RNAse-free water. Bulk mRNA-seq with polyA selection and 30 million reads/samples was performed by Genewiz, Azenta Life sciences in Oxford, using Illumina, 2×150b system. All samples passed QC.

Quality control was performed using FastQC/MultiQC, assessing the yield, number and percentage of duplicate reads, the per sequence Phred quality score, the GC content, the length distribution, and overrepresenting sequences or adapter contaminations. Reads were mapped to the GRCh38 using the STAR aligner with ENCODE options, and gene counts were determined using HTSeq. DESeq2 was used to determine differential gene expression. Moderation of log fold changes was carried out by using a zero-mean normal prior on the coefficients of interest. Gene ontology (GO) analyses were performed in R using the GOseq package.

In order to compile the characteristic gene signatures for human DRGs we calculated the 1st principal component - eigengene of the variance stabilized expression values for the neuronal genes expressed in DRG’s from human organ donors (P. Ray et al., 2018).

Single-cell RNA-seq from human and mouse DRGs was used to predict cell type proportions in bulk RNA-seq by doing deconvolution analysis with the SCDC R package (Bhuiyan et al., 2024). Full methodology in **Supplemental Methods**.

### Electrophysiological recordings

Whole-cell patch clamp recordings of HD10.6 cells were performed at 21 days following maturation (3 separate maturations), using an Axopatch 200B amplifier and Digidata 1440 acquisition system (Molecular Devices). Data were low-passed filtered at 2 kHz and sampled at 10-20 kHz. Series resistance was compensated 70-85% to reduce voltage errors. All data were analysed in Clampfit 10 software.

For current-clamp recordings, borosilicate glass capillaries (1.5 mm OD, 0.84 mm ID; World Precision Instruments) were pulled to form patch pipettes of 2–5 MΩ tip resistance and filled with an internal solution containing (mM): 100 K-gluconate, 28 KCl, 1 MgCl2, 5 Mg-ATP, 10 HEPES, and 0.5 EGTA; pH was adjusted to 7.3 with KOH and osmolarity set at 305-310 mOsm. Extracellular solution contained (mM): 140 NaCl, 4.7 KCl, 2.5 CaCl2, 1.2 MgCl2, 10 HEPES and 10 glucose; pH was adjusted to 7.4 with NaOH and osmolarity set at 310-315 mOsm. Liquid junction potential (∼13 mV) was corrected. Input resistance (Rin) was derived by measuring the membrane deflection caused by 3 hyperpolarising current steps (-30, -20 and -10 pA). Rheobase was determined by applying 50 ms depolarising currents of increasing steps (10 pA) until action potential (AP) generation. Repetitive firing was assessed by applying prolonged (500 ms) depolarising currents of increasing steps (20 pA). Resting membrane potential (RMP) was measured in bridge mode (I=0). Only cells with a RMP greater than -45 mV were considered for analysis.

For voltage-clamp recordings, patch pipettes of 1–2 MΩ tip were filled with an internal solution containing (mM): 140 CsF, 10 NaCl, 2 MgCl2, 0.1 CaCl2, 10 HEPES, and 1.1 EGTA; pH was adjusted to 7.3 with KOH and osmolarity set at 305-310 mOsm. Extracellular solution contained (mM): 70 NaCl, 50 Choline chloride, 20 Tetraethylammonium chloride, 3 KCl, 1 CaCl2, 1 MgCl2, 10 HEPES and 10 glucose; pH was adjusted to 7.4 with NaOH and osmolarity set at 310-315 mOsm. To assess voltage-gated Na+ currents, membrane potential was stepped from -80 mV to +40 mV in 10 mV increments, from a holding potential of -100 mV. TTX (200 nM) was added to the extracellular solution to measure TTX-resistant currents. Recordings were discarded if series resistance was greater than 10 MΩ or deviated by more than 20% during the recording. Linear leak subtraction was performed during acquisition using P/4 leak subtraction. The peak amplitude at each step was used to calculate the I-V curve. Current density was estimated using the maximum peak amplitude.

### Fura-2 AM Calcium Imaging

For Fura-2 AM Imaging, HD10.6 neurons cultured and matured to relevant days on coated-coverslips and were loaded with 5 µM Fura-2 AM (Invitrogen F1221) for 30 min culture medium. Cells were washed once with extracellular solution (ECS) containing (mM): NaCl 140, KCl 5, CaCl_2_ 2 MgCl2 1, HEPES 10 and Glucose 10; pH was adjusted to 7.3 with NaOH and osmolarity set at 310-315 mOsm. ECS containing PGE2 (1µM), α,β-MeATP (100 µM), AITC (50µM), capsaicin (1 µM), menthol (100 µM) or KCl solution (50 mM) were applied for 30 s followed by a 60 - 120 s ECS wash. Cells were imaged at 1 Hz under continuous perfusion (Bioscience tools perfusion system and software) using a Zeiss Axio Observer inverted microscope and HBO 100 Microscope illuminating system.

### GCaMP8f calcium imaging

For GCaMP Imaging, HD10.6 sensory neurons plated and matured to DIV7 in microfluidic devices and transduced with AAV-CAG-jGCaMP8f-WPRE (Addgene #179254; 500 K MOI/cell) for 7 days in maturation medium, after which media fully replaced. After 21 days transduction, cells were imaged with 20X objective using a spinning-disk confocal microscope (IXplore IX83 SpinSR Super-Resolution System), at 10 Hz for 30 seconds, to determine baseline, and imaged again immediately after treatment of 1µM Prostaglandin E2.

For analysis, fluorescence activity was expressed as ΔF/F₀, where F₀ corresponded to the 8th percentile of baseline fluorescence. Integrated fluorescence in which sum of the area under the curve of ΔF/F₀ traces was used to quantify calcium transient activity.

### Lentiviral Transduction (pLenti-Synapsin-EYFP-WPRE)

HD10.6 cells were transduced with the lentiviral construct pLenti-Synapsin-EYFP-WPRE (Steinbeck et al., 2016) (gift from Dr Lucy Farrimond, University of Oxford, UK) at a multiplicity of infection (MOI) of 10 for 7 days under standard culture conditions. This construct drives enhanced EYFP expression, enabling neuronal labelling. Following transduction, EYFP-expressing HD10.6 cells were used for live-cell imaging in neurite regeneration assays.

### Neurite Regeneration Assay in HD10.6 Human DRG-Derived Neurons Using Microfluidic Devices

Microfluidic devices were created in-house using epoxy templates as previously described (Clark et al., 2018) and mounted onto glass-bottom dishes (WillCo). Devices were plasma-treated (Femto plasma cleaner; Diener Electronics, Germany) prior to coating with Cultrex extracellular matrix. EYFP-expressing HD10.6 cells were seeded into the soma compartment in proliferation medium. Twenty-four hours after seeding (designated day 0), neuronal differentiation was initiated by replacing proliferation medium with maturation medium containing either neurotrophic factors (NGF, 50 ng/mL; GDNF, 25 ng/mL) or control conditions lacking trophic support. Half-volume media changes were performed every 2–4 days, and cultures were maintained for up to 21 days in vitro.

On day 20 in vitro, live-cell imaging was performed using a spinning-disk confocal microscope (IXplore IX83 SpinSR Super-Resolution System) by acquiring four fields of view from predefined locations within the axonal compartment while maintaining cells at 37 °C and 5% CO₂. On day 21, axotomy was performed by vacuum aspiration restricted to the axonal compartment. Twenty-four hours after injury, the same fields were re-imaged to assess neurite regeneration. Regeneration was quantified by measuring the area occupied by EYFP-positive neurites and normalised to pre-axotomy values using ImageJ software.

### Vacor and DSRM-3716 treatment viability and fragmentation assays

HD10.6 cells were seeded onto poly-D-lysine- and Cultrex-coated plates and maintained in proliferation medium before induction of maturation. Cells were matured to DIV28 in maturation medium prior to drug treatment.

For ATP viability assays, DIV28 HD10.6 cells were cultured in white opaque 96-well plates (Greiner) and treated in full maturation medium with DMSO vehicle or vacor (Greyhound Chromatography N-13738) at 1, 10, 25, or 50 µM. Vehicle control wells received 0.1% DMSO, matched to the highest DMSO concentration used in vacor-treated wells. Cell viability was measured at 24, 48, and 72 h using the CellTiter-Glo® Luminescent Cell Viability Assay according to the manufacturer’s instructions. Luminescence values were background-subtracted, normalised to DMSO-treated controls at each timepoint, and used to generate dose–response curves for IC50 estimation.

For axonal fragmentation assays, HD10.6 cells were seeded onto poly-D-lysine- and Cultrex-coated black imaging 24-well or 96-well plates (Ibidi) and matured as above. At DIV7, cells were transduced with pAAV-CAG-tdTomato (Addgene #59462) at an MOI of 100,000 for 7 days to label neuronal processes. At DIV14, rat Schwann cells were added to generate co-cultures under pro-myelinating conditions adapted from Clark et al. (Clark et al., 2017) At DIV28, co-cultures were treated with DMSO vehicle or vacor at 1, 10, 25, or 50 µM for 6, 24, or 48 h. Vehicle control wells received 0.1% DMSO. Plates were live-imaged using a spinning-disk confocal microscope (IXplore IX83 SpinSR Super-Resolution System), with 8–20 images acquired per well depending on plate format. tdTomato-positive axons were imaged using consistent acquisition settings across treatment groups, and axonal degeneration was quantified by fragmentation index analysis.

To assess pharmacological rescue, DIV28 tdTomato-labelled HD10.6 co-cultures were treated with 25 µM vacor alone or 25 µM vacor in combination with DSRM-3716 at 0.1, 0.3, 1, 3, or 10 µM. Vehicle control wells received 0.11% DMSO, matched to the highest DMSO concentration used in the combined treatment conditions. Cell viability and axonal fragmentation were quantified as described above. Fragmentation index was calculated from thresholded tdTomato-positive axonal masks as the proportion of fragmented axonal signal relative to total axonal signal. Full image-processing and fragmentation index calculation details are provided in Supplementary Figure 8.

### Microscopy

Fluorescence microscopy images were acquired on a spinning-disk confocal microscope (IXplore IX83 SpinSR Super-Resolution System) using 20X or 40X objectives.

### Image Analysis

Segmentation and fluorescence quantification were performed via Cellpose (cyto3 model) and ImageJ (FIJI). ROIs corresponding to individual cells were extracted from maximum intensity projections and applied to time-lapse sequences to measure mean fluorescence intensity over time. Traces were background-subtracted, normalised to baseline (ΔF/F₀), and exported to Python for quantitative analysis.

### Statistical Analysis

All statistical analyses were performed using GraphPad Prism 10. Data normality was assessed using the Shapiro-Wilk test. Non-parametric data were analysed using the Kruskal-Wallis test followed by Dunn’s multiple-comparisons post hoc test, or the paired Wilcoxon signed-rank test, where appropriate. Parametric data were analysed using t-tests or one-way ANOVA followed by Tukey’s multiple-comparisons post hoc test, where appropriate. Data are presented as mean ± SEM, unless stated otherwise. Statistical significance was defined as P < 0.05.

## 5. Acknowledgements

We thank Prof. Derek C. Molliver (University of New England, Biddeford, ME, USA) for kindly providing the HD10.6 cell line and for fruitful discussions on this cell line.

This work was funded by: Wellcome Trust (Wellcome Investigator Grant to DB, 223149/Z/21/Z, Wellcome collaborative awards: 220906/Z/20/Z and 224257/Z/21/Z) and the UK Medical Research Council (grant ref. MR/T020113/1).

## References

Akopian, A. N., Sivilotti, L., & Wood, J. N. (1996). A tetrodotoxin-resistant voltage-gated sodium channel expressed by sensory neurons. Nature, 379(6562), 257–262. 10.1038/379257a0

Al-Abbasi, Z., Bhuiyan, S. A., Renthal, W., & Molliver, D. C. (2025). A Transcriptomic Comparison of the HD10.6 Human Sensory Neuron-Derived Cell Line with Primary and iPSC Sensory Neurons. bioRxiv: The Preprint Server for Biology, 2025.04.03.643725. 10.1101/2025.04.03.643725

Barry, A. M., Zhao, N., Yang, X., Bennett, D. L., & Baskozos, G. (2023). Deep RNA-seq of male and female murine sensory neuron subtypes after nerve injury. PAIN, 164(10), 2196. 10.1097/j.pain.0000000000002934

Baskozos, G., Hébert, H. L., Pascal, M. M., Themistocleous, A. C., Macfarlane, G. J., Wynick, D., Bennett, D. L., & Smith, B. H. (2023). Epidemiology of neuropathic pain: An analysis of prevalence and associated factors in UK Biobank. Pain Reports, 8(2), e1066. 10.1097/PR9.0000000000001066

Ben-David, U., Siranosian, B., Ha, G., Tang, H., Oren, Y., Hinohara, K., Strathdee, C. A., Dempster, J., Lyons, N. J., Burns, R., Nag, A., Kugener, G., Cimini, B., Tsvetkov, P., Maruvka, Y. E., O’Rourke, R., Garrity, A., Tubelli, A. A., Bandopadhayay, P., … Golub, T. R. (2018). Genetic and transcriptional evolution alters cancer cell line drug response. Nature, 560(7718), 325–330. 10.1038/s41586-018-0409-3

Bennett, D. L., Clark, A. J., Huang, J., Waxman, S. G., & Dib-Hajj, S. D. (2019). The Role of Voltage-Gated Sodium Channels in Pain Signaling. Physiological Reviews, 99(2), 1079–1151. 10.1152/physrev.00052.2017

Bhuiyan, S. A., Nagi, S. S., Sankaranarayanan, I., Semizoglou, E., Usoskin, D., Yang, L., Yu, H., Arendt-Tranholm, A., Bertels, Z., Bhatia, P., Bouchatta, O., Boyer, K., Cervantes, A., Chalif, J., Chintalapudi, H., Cicalo, A., Copits, B., Cronin, C., Curatolo, M., … Network, N. P. H. P. (2025). A Reference Atlas of the Human Dorsal Root Ganglion (p. 2025.11.05.686654). bioRxiv. 10.1101/2025.11.05.686654

Bhuiyan, S. A., Xu, M., Yang, L., Semizoglou, E., Bhatia, P., Pantaleo, K. I., Tochitsky, I., Jain, A., Erdogan, B., Blair, S., Cat, V., Mwirigi, J. M., Sankaranarayanan, I., Tavares-Ferreira, D., Green, U., McIlvried, L. A., Copits, B. A., Bertels, Z., Del Rosario, J. S., … Renthal, W. (2024). Harmonized cross-species cell atlases of trigeminal and dorsal root ganglia. Science Advances, 10(25), eadj9173. 10.1126/sciadv.adj9173

Bosanac, T., Hughes, R. O., Engber, T., Devraj, R., Brearley, A., Danker, K., Young, K., Kopatz, J., Hermann, M., Berthemy, A., Boyce, S., Bentley, J., & Krauss, R. (2021). Pharmacological SARM1 inhibition protects axon structure and function in paclitaxel-induced peripheral neuropathy. Brain, 144(10), 3226–3238. 10.1093/brain/awab184

Breivik, H., Collett, B., Ventafridda, V., Cohen, R., & Gallacher, D. (2006). Survey of chronic pain in Europe: Prevalence, impact on daily life, and treatment. European Journal of Pain, 10(4), 287–333. 10.1016/j.ejpain.2005.06.009

Brinkmann, B. G., Agarwal, A., Sereda, M. W., Garratt, A. N., Müller, T., Wende, H., Stassart, R. M., Nawaz, S., Humml, C., Velanac, V., Radyuschkin, K., Goebbels, S., Fischer, T. M., Franklin, R. J., Lai, C., Ehrenreich, H., Birchmeier, C., Schwab, M. H., & Nave, K.-A. (2008). Neuregulin-1/ErbB signaling serves distinct functions in myelination of the peripheral and central nervous system. Neuron, 59(4), 581–595. 10.1016/j.neuron.2008.06.028

Cao, L., McDonnell, A., Nitzsche, A., Alexandrou, A., Saintot, P.-P., Loucif, A. J. C., Brown, A. R., Young, G., Mis, M., Randall, A., Waxman, S. G., Stanley, P., Kirby, S., Tarabar, S., Gutteridge, A., Butt, R., McKernan, R. M., Whiting, P., Ali, Z., … Stevens, E. B. (2016). Pharmacological reversal of a pain phenotype in iPSC-derived sensory neurons and patients with inherited erythromelalgia. Science Translational Medicine, 8(335), 335ra56. 10.1126/scitranslmed.aad7653

Chambers, S. M., Qi, Y., Mica, Y., Lee, G., Zhang, X.-J., Niu, L., Bilsland, J., Cao, L., Stevens, E., Whiting, P., Shi, S.-H., & Studer, L. (2012). Combined small-molecule inhibition accelerates developmental timing and converts human pluripotent stem cells into nociceptors. Nature Biotechnology, 30(7), 715–720. 10.1038/nbt.2249

Chen, S., Velardez, M. O., Warot, X., Yu, Z.-X., Miller, S. J., Cros, D., & Corfas, G. (2006). Neuregulin 1–erbB Signaling Is Necessary for Normal Myelination and Sensory Function. The Journal of Neuroscience, 26(12), 3079–3086. 10.1523/JNEUROSCI.3785-05.2006

Chen, Y.-C., Bullock, B., Ogbeh, D. I., Ai, J., Okoh, J., Liu, J., Qiang, Y., & Hsia, S. V. (2026). In vitro modeling of human dorsal root ganglion neurons for GCaMP6-based calcium imaging of sensory responses to HSV-1 infection. Journal of Virological Methods, 339, 115264. 10.1016/j.jviromet.2025.115264

Clark, A. J., Kaller, M. S., Galino, J., Willison, H. J., Rinaldi, S., & Bennett, D. L. H. (2017). Co-cultures with stem cell-derived human sensory neurons reveal regulators of peripheral myelination. Brain, 140(4), 898–913. 10.1093/brain/awx012

Clark, A. J., Kugathasan, U., Baskozos, G., Priestman, D. A., Fugger, N., Lone, M. A., Othman, A., Chu, K. H., Blesneac, I., Wilson, E. R., Laurà, M., Kalmar, B., Greensmith, L., Hornemann, T., Platt, F. M., Reilly, M. M., & Bennett, D. L. (2021). An iPSC model of hereditary sensory neuropathy-1 reveals L-serine-responsive deficits in neuronal ganglioside composition and axoglial interactions. *Cell Reports*. Medicine, 2(7), 100345. 10.1016/j.xcrm.2021.100345

Clark, A. J., Menendez, G., AlQatari, M., Patel, N., Arstad, E., Schiavo, G., & Koltzenburg, M. (2018). Functional imaging in microfluidic chambers reveals sensory neuron sensitivity is differentially regulated between neuronal regions. Pain, 159(7), 1413–1425. 10.1097/j.pain.0000000000001145

Cummins, T. R., Dib-Hajj, S. D., Black, J. A., Akopian, A. N., Wood, J. N., & Waxman, S. G. (1999). A novel persistent tetrodotoxin-resistant sodium current in SNS-null and wild-type small primary sensory neurons. The Journal of Neuroscience: The Official Journal of the Society for Neuroscience, 19(24), RC43. 10.1523/JNEUROSCI.19-24-j0001.1999

Deng, T., Jovanovic, V. M., Tristan, C. A., Weber, C., Chu, P.-H., Inman, J., Ryu, S., Jethmalani, Y., Ferreira de Sousa, J., Ormanoglu, P., Twumasi, P., Sen, C., Shim, J., Jayakar, S., Bear Zhang, H.-X., Jo, S., Yu, W., Voss, T. C., Simeonov, A., … Singeç, I. (2023). Scalable generation of sensory neurons from human pluripotent stem cells. Stem Cell Reports, 18(4), 1030–1047. 10.1016/j.stemcr.2023.03.006

Dhaka, A., Earley, T. J., Watson, J., & Patapoutian, A. (2008). Visualizing Cold Spots: TRPM8-Expressing Sensory Neurons and Their Projections. The Journal of Neuroscience, 28(3), 566–575. 10.1523/JNEUROSCI.3976-07.2008

Diaz, P., Awadelkareem, M. A., Muñoz, D., Espinoza, F., Clark, A. J., Altermatt, F., Veliz, L., Mora, A. S., Nyström, A., Fuentes, I., Bennett, D., & Calvo, M. (2026). Impaired skin reinnervation in Epidermolysis Bullosa due to neurotrophic deficiency. Pain. 10.1097/j.pain.0000000000004001

Dib-Hajj, S., Black, J. A., Cummins, T. R., & Waxman, S. G. (2002). NaN/Nav1.9: A sodium channel with unique properties. Trends in Neurosciences, 25(5), 253–259. 10.1016/s0166-2236(02)02150-1

Dochnal, S. A., Du, Y., Bandari, D., Franco Malange, K., Bryant, J., Lemes, J. B. P., Whitford, A., Cliffe, A. R., Mali, P., Dore, K., Miller, Y. I., & Yaksh, T. L. (2026). Human Dorsal Root Ganglia Neuronal Cell Line to Study Nociceptive Signaling: A New Pipeline for Pain Therapy. The FASEB Journal, 40(3), e71528. 10.1096/fj.202503698R

Dochnal, S. A., Du, Y., Bandari, D., Malange, K. F., Bryant, J., Lemes, J. B. P., Whitford, A., Cliffe, A. R., Mali, P., Dore, K., Miller, Y. I., & Yaksh, T. L. (2025). Human dorsal root ganglia neuronal cell line to study nociceptive signaling: A new pipeline for pain therapy (p. 2025.09.27.674468). bioRxiv. 10.1101/2025.09.27.674468

Dubin, A. E., & Patapoutian, A. (2010). Nociceptors: The sensors of the pain pathway. The Journal of Clinical Investigation, 120(11), 3760–3772. 10.1172/JCI42843

Dykes, I. M., Lanier, J., Raisa Eng, S., & Turner, E. E. (2010). Brn3a regulates neuronal subtype specification in the trigeminal ganglion by promoting Runx expression during sensory differentiation. Neural Development, 5(1), 3. 10.1186/1749-8104-5-3

Eberhardt, E., Havlicek, S., Schmidt, D., Link, A. S., Neacsu, C., Kohl, Z., Hampl, M., Kist, A. M., Klinger, A., Nau, C., Schüttler, J., Alzheimer, C., Winkler, J., Namer, B., Winner, B., & Lampert, A. (2015). Pattern of Functional TTX-Resistant Sodium Channels Reveals a Developmental Stage of Human iPSC- and ESC-Derived Nociceptors. Stem Cell Reports, 5(3), 305–313. 10.1016/j.stemcr.2015.07.010

Essuman, K., Summers, D. W., Sasaki, Y., Mao, X., DiAntonio, A., & Milbrandt, J. (2017). The SARM1 Toll/Interleukin-1 Receptor Domain Possesses Intrinsic NAD+ Cleavage Activity that Promotes Pathological Axonal Degeneration. Neuron, 93(6), 1334–1343.e5. 10.1016/j.neuron.2017.02.022

Faber, C. G., Lauria, G., Merkies, I. S. J., Cheng, X., Han, C., Ahn, H.-S., Persson, A.-K., Hoeijmakers, J. G. J., Gerrits, M. M., Pierro, T., Lombardi, R., Kapetis, D., Dib-Hajj, S. D., & Waxman, S. G. (2012). Gain-of-function Na_v_ 1.8 mutations in painful neuropathy. Proceedings of the National Academy of Sciences, 109(47), 19444–19449. 10.1073/pnas.1216080109

Figley, M. D., Gu, W., Nanson, J. D., Shi, Y., Sasaki, Y., Cunnea, K., Malde, A. K., Jia, X., Luo, Z., Saikot, F. K., Mosaiab, T., Masic, V., Holt, S., Hartley-Tassell, L., McGuinness, H. Y., Manik, M. K., Bosanac, T., Landsberg, M. J., Kerry, P. S., … Ve, T. (2021). SARM1 is a metabolic sensor activated by an increased NMN/NAD+ ratio to trigger axon degeneration. Neuron, 109(7), 1118–1136.e11. 10.1016/j.neuron.2021.02.009

Fofie, C. K., Granja-Vazquez, R., Truong, V., Walsh, P., Price, T., Biswas, S., Dussor, G., Pancrazio, J., & Kolber, B. (2025). Profiling human iPSC-derived sensory neurons for analgesic drug screening using a multi-electrode array. Cell Reports Methods, 5(5), 101051. 10.1016/j.crmeth.2025.101051

Geisler, S., Huang, S. X., Strickland, A., Doan, R. A., Summers, D. W., Mao, X., Park, J., DiAntonio, A., & Milbrandt, J. (2019). Gene therapy targeting SARM1 blocks pathological axon degeneration in mice. Journal of Experimental Medicine, 216(2), 294–303. 10.1084/jem.20181040

Ghislain, J., & Charnay, P. (2006). Control of myelination in Schwann cells: A Krox20 cis-regulatory element integrates Oct6, Brn2 and Sox10 activities. EMBO Reports, 7(1), 52–58. 10.1038/sj.embor.7400573

Gilley, J., Orsomando, G., Nascimento-Ferreira, I., & Coleman, M. P. (2015). Absence of SARM1 Rescues Development and Survival of NMNAT2-Deficient Axons. Cell Reports, 10(12), 1974–1981. 10.1016/j.celrep.2015.02.060

Haberberger, R. V., Barry, C., & Matusica, D. (2020). Immortalized Dorsal Root Ganglion Neuron Cell Lines. Frontiers in Cellular Neuroscience, 14. 10.3389/fncel.2020.00184

Han, C., Estacion, M., Huang, J., Vasylyev, D., Zhao, P., Dib-Hajj, S. D., & Waxman, S. G. (2015). Human Na(v)1.8: Enhanced persistent and ramp currents contribute to distinct firing properties of human DRG neurons. Journal of Neurophysiology, 113(9), 3172–3185. 10.1152/jn.00113.2015

Hjerling-Leffler, J., AlQatari, M., Ernfors, P., & Koltzenburg, M. (2007). Emergence of Functional Sensory Subtypes as Defined by Transient Receptor Potential Channel Expression. The Journal of Neuroscience, 27(10), 2435–2443. 10.1523/JNEUROSCI.5614-06.2007

Kalia, A. K., Rösseler, C., Granja-Vazquez, R., Ahmad, A., Pancrazio, J. J., Neureiter, A., Zhang, M., Sauter, D., Vetter, I., Andersson, A., Dussor, G., Price, T. J., Kolber, B. J., Truong, V., Walsh, P., & Lampert, A. (2024). How to differentiate induced pluripotent stem cells into sensory neurons for disease modelling: A functional assessment. Stem Cell Research & Therapy, 15(1), 99. 10.1186/s13287-024-03696-2

Karashima, Y., Damann, N., Prenen, J., Talavera, K., Segal, A., Voets, T., & Nilius, B. (2007). Bimodal Action of Menthol on the Transient Receptor Potential Channel TRPA1. The Journal of Neuroscience, 27(37), 9874–9884. 10.1523/JNEUROSCI.2221-07.2007

Kim, J.-I., Imaizumi, K., Jurjuț, O., Kelley, K. W., Wang, D., Thete, M. V., Hudacova, Z., Amin, N. D., Levy, R. J., Scherrer, G., & Pașca, S. P. (2025). Human assembloid model of the ascending neural sensory pathway. Nature, 642(8066), 143–153. 10.1038/s41586-025-08808-3

Lampert, A., Bennett, D. L., McDermott, L. A., Neureiter, A., Eberhardt, E., Winner, B., & Zenke, M. (2020). Human sensory neurons derived from pluripotent stem cells for disease modelling and personalized medicine. Neurobiology of Pain, 8, 100055. 10.1016/j.ynpai.2020.100055

LeBlang, C. J., Pazyra-Murphy, M. F., Silagi, E. S., Dasgupta, S., Tsolias, M., Miller, T., Petrova, V., Zhen, S., Jovanovic, V. M., Castellano, D., Gerrish, K., Ormanoglu, P., Tristan, C. A., Singeç, I., Woolf, C. J., Tasdemir-Yilmaz, O., & Segal, R. A. (2025). Satellite glial contact enhances differentiation and maturation of human iPSC-derived sensory neurons. Stem Cell Reports, 20(10), 102639. 10.1016/j.stemcr.2025.102639

Liang, C., Tao, Y., Shen, C., Tan, Z., Xiong, W.-C., & Mei, L. (2012). Erbin Is Required for Myelination in Regenerated Axons after Injury. Journal of Neuroscience, 32(43), 15169–15180. 10.1523/JNEUROSCI.2466-12.2012

Loreto, A., Angeletti, C., Gu, W., Osborne, A., Nieuwenhuis, B., Gilley, J., Merlini, E., Arthur-Farraj, P., Amici, A., Luo, Z., Hartley-Tassell, L., Ve, T., Desrochers, L. M., Wang, Q., Kobe, B., Orsomando, G., & Coleman, M. P. (2021). Neurotoxin-mediated potent activation of the axon degeneration regulator SARM1. eLife, 10, e72823. 10.7554/eLife.72823

Lund, R. J., Nikula, T., Rahkonen, N., Närvä, E., Baker, D., Harrison, N., Andrews, P., Otonkoski, T., & Lahesmaa, R. (2012). High-throughput karyotyping of human pluripotent stem cells. Stem Cell Research, 9(3), 192–195. 10.1016/j.scr.2012.06.008

Mani, A., Mendel, M., Westwood, P., Bonardi, C., Saal, W., Topp, A., Bilyard, M., Brigo, A., Wittwer, M. B., Benz, J., Kuhn, B., Haider, A., Giroud, M., & Keaney, J. (2025). SARM1 base-exchange inhibitors induce SARM1 activation and neurodegeneration at low doses. Npj Drug Discovery, 2(1), 12. 10.1038/s44386-025-00016-3

Mazzara, P. G., Muggeo, S., Luoni, M., Massimino, L., Zaghi, M., Valverde, P. T.-T., Brusco, S., Marzi, M. J., Palma, C., Colasante, G., Iannielli, A., Paulis, M., Cordiglieri, C., Giannelli, S. G., Podini, P., Gellera, C., Taroni, F., Nicassio, F., Rasponi, M., & Broccoli, V. (2020). Frataxin gene editing rescues Friedreich’s ataxia pathology in dorsal root ganglia organoid-derived sensory neurons. Nature Communications, 11(1), 4178. 10.1038/s41467-020-17954-3

Middleton, S. J., Barry, A. M., Comini, M., Li, Y., Ray, P. R., Shiers, S., Themistocleous, A. C., Uhelski, M. L., Yang, X., Dougherty, P. M., Price, T. J., & Bennett, D. L. (2021). Studying human nociceptors: From fundamentals to clinic. Brain, 144(5), 1312–1335. 10.1093/brain/awab048

Moparthi, L., Sinica, V., Moparthi, V. K., Kreir, M., Vignane, T., Filipovic, M. R., Vlachova, V., & Zygmunt, P. M. (2022). The human TRPA1 intrinsic cold and heat sensitivity involves separate channel structures beyond the N-ARD domain. Nature Communications, 13(1), 6113. 10.1038/s41467-022-33876-8

Park, J. W., Bae, S. J., Yun, J. H., Kim, S., & Park, M. (2024). Assessment of Genetic Stability in Human Induced Pluripotent Stem Cell-Derived Cardiomyocytes by Using Droplet Digital PCR. International Journal of Molecular Sciences, 25(2), 1101. 10.3390/ijms25021101

Ray, P. R., Shiers, S., Caruso, J. P., Tavares-Ferreira, D., Sankaranarayanan, I., Uhelski, M. L., Li, Y., North, R. Y., Tatsui, C., Dussor, G., Burton, M. D., Dougherty, P. M., & Price, T. J. (2023). RNA profiling of human dorsal root ganglia reveals sex differences in mechanisms promoting neuropathic pain. Brain: A Journal of Neurology, 146(2), 749–766. 10.1093/brain/awac266

Ray, P., Torck, A., Quigley, L., Wangzhou, A., Neiman, M., Rao, C., Lam, T., Kim, J.-Y., Kim, T. H., Zhang, M. Q., Dussor, G., & Price, T. J. (2018). Comparative transcriptome profiling of the human and mouse dorsal root ganglia: An RNA-seq-based resource for pain and sensory neuroscience research. Pain, 159(7), 1325–1345. 10.1097/j.pain.0000000000001217

Raymon, H. K., Thode, S., Zhou, J., Friedman, G. C., Pardinas, J. R., Barrere, C., Johnson, R. M., & Sah, D. W. Y. (1999). Immortalized Human Dorsal Root Ganglion Cells Differentiate into Neurons with Nociceptive Properties. The Journal of Neuroscience, 19(13), 5420–5428. 10.1523/JNEUROSCI.19-13-05420.1999

Ren, L., Chang, M. J., Zhang, Z., Dhaka, A., Guo, Z., & Cao, Y.-Q. (2018). Quantitative analysis of mouse dural afferent neurons expressing TRPM8, VGLUT3 and NF200. Headache, 58(1), 88–101. 10.1111/head.13188

Ruscheweyh, R., Forsthuber, L., Schoffnegger, D., & Sandkühler, J. (2007). Modification of classical neurochemical markers in identified primary afferent neurons with Abeta-, Adelta-, and C-fibers after chronic constriction injury in mice. The Journal of Comparative Neurology, 502(2), 325–336. 10.1002/cne.21311

Schinke, C., Fernandez Vallone, V., Ivanov, A., Peng, Y., Körtvelyessy, P., Nolte, L., Huehnchen, P., Beule, D., Stachelscheid, H., Boehmerle, W., & Endres, M. (2021). Modeling chemotherapy induced neurotoxicity with human induced pluripotent stem cell (iPSC) -derived sensory neurons. Neurobiology of Disease, 155, 105391. 10.1016/j.nbd.2021.105391

Schrenk-Siemens, K., Wende, H., Prato, V., Song, K., Rostock, C., Loewer, A., Utikal, J., Lewin, G. R., Lechner, S. G., & Siemens, J. (2015). PIEZO2 is required for mechanotransduction in human stem cell–derived touch receptors. Nature Neuroscience, 18(1), 10–16. 10.1038/nn.3894

Schwartzentruber, J., Foskolou, S., Kilpinen, H., Rodrigues, J., Alasoo, K., Knights, A. J., Patel, M., Goncalves, A., Ferreira, R., Benn, C. L., Wilbrey, A., Bictash, M., Impey, E., Cao, L., Lainez, S., Loucif, A. J., Whiting, P. J., HIPSCI Consortium, Gutteridge, A., & Gaffney, D. J. (2018). Molecular and functional variation in iPSC-derived sensory neurons. Nature Genetics, 50(1), 54–61. 10.1038/s41588-017-0005-8

Seamon, K. B., Padgett, W., & Daly, J. W. (1981). Forskolin: Unique diterpene activator of adenylate cyclase in membranes and in intact cells. Proceedings of the National Academy of Sciences, 78(6), 3363–3367. 10.1073/pnas.78.6.3363

Steichen, C., Hannoun, Z., Luce, E., Hauet, T., & Dubart-Kupperschmitt, A. (2019). Genomic integrity of human induced pluripotent stem cells: Reprogramming, differentiation and applications. World Journal of Stem Cells, 11(10), 729–747. 10.4252/wjsc.v11.i10.729

Steinbeck, J. A., Jaiswal, M. K., Calder, E. L., Kishinevsky, S., Weishaupt, A., Toyka, K. V., Goldstein, P. A., & Studer, L. (2016). Functional Connectivity under Optogenetic Control Allows Modeling of Human Neuromuscular Disease. Cell Stem Cell, 18(1), 134–143. 10.1016/j.stem.2015.10.002

Story, G. M., Peier, A. M., Reeve, A. J., Eid, S. R., Mosbacher, J., Hricik, T. R., Earley, T. J., Hergarden, A. C., Andersson, D. A., Hwang, S. W., McIntyre, P., Jegla, T., Bevan, S., & Patapoutian, A. (2003). ANKTM1, a TRP-like Channel Expressed in Nociceptive Neurons, Is Activated by Cold Temperatures. Cell, 112(6), 819–829. 10.1016/S0092-8674(03)00158-2

Tavares-Ferreira, D., Shiers, S., Ray, P. R., Wangzhou, A., Jeevakumar, V., Sankaranarayanan, I., Cervantes, A. M., Reese, J. C., Chamessian, A., Copits, B. A., Dougherty, P. M., Gereau, R. W., Burton, M. D., Dussor, G., & Price, T. J. (2022). Spatial transcriptomics of dorsal root ganglia identifies molecular signatures of human nociceptors. Science Translational Medicine, 14(632), eabj8186. 10.1126/scitranslmed.abj8186

Topilko, P., Schneider-Maunoury, S., Levi, G., Baron-Van Evercooren, A., Chennoufi, A. B. Y., Seitanidou, T., Babinet, C., & Charnay, P. (1994). Krox-20 controls myelination in the peripheral nervous system. Nature, 371(6500), 796–799. 10.1038/371796a0

Van Lent, J., Prior, R., Pérez Siles, G., Cutrupi, A. N., Kennerson, M. L., Vangansewinkel, T., Wolfs, E., Mukherjee-Clavin, B., Nevin, Z., Judge, L., Conklin, B., Tyynismaa, H., Clark, A. J., Bennett, D. L., Van Den Bosch, L., Saporta, M., & Timmerman, V. (2024). Advances and challenges in modeling inherited peripheral neuropathies using iPSCs. Experimental & Molecular Medicine, 56(6), 1348–1364. 10.1038/s12276-024-01250-x

Xiong, C., Chua, K. C., Stage, T. B., Priotti, J., Kim, J., Altman-Merino, A., Chan, D., Saraf, K., Canato Ferracini, A., Fattahi, F., & Kroetz, D. L. (2021). Human Induced Pluripotent Stem Cell Derived Sensory Neurons are Sensitive to the Neurotoxic Effects of Paclitaxel. Clinical and Translational Science, 14(2), 568–581. 10.1111/cts.12912

Yu, H., Nagi, S. S., Usoskin, D., Hu, Y., Kupari, J., Bouchatta, O., Yan, H., Cranfill, S. L., Gautam, M., Su, Y., Lu, Y., Wymer, J., Glanz, M., Albrecht, P., Song, H., Ming, G.-L., Prouty, S., Seykora, J., Wu, H., … Luo, W. (2024). Leveraging deep single-soma RNA sequencing to explore the neural basis of human somatosensation. Nature Neuroscience, 27(12), 2326–2340. 10.1038/s41593-024-01794-1

Zhang, Q., Martin-Caraballo, M., & Hsia, S. V. (2020). Modulation of Voltage-Gated Sodium Channel Activity in Human Dorsal Root Ganglion Neurons by Herpesvirus Quiescent Infection. Journal of Virology, 94(3), e01823–19. 10.1128/JVI.01823-19

Zhang, Y., Rózsa, M., Liang, Y., Bushey, D., Wei, Z., Zheng, J., Reep, D., Broussard, G. J., Tsang, A., Tsegaye, G., Narayan, S., Obara, C. J., Lim, J.-X., Patel, R., Zhang, R., Ahrens, M. B., Turner, G. C., Wang, S. S.-H., Korff, W. L., … Looger, L. L. (2023). Fast and sensitive GCaMP calcium indicators for imaging neural populations. Nature, 615(7954), 884–891. 10.1038/s41586-023-05828-9

