## Supplementary Methods for "Extended maturation of the HD10.6 immortalised human dorsal root ganglion cell line enables modelling of nociceptive responses and neural injury"

**RNA-seq**

Quality control was performed using FastQC/MultiQC, assessing the yield, number and percentage of duplicate reads, the per sequence Phred quality score, the GC content, the length distribution, and overrepresenting sequences or adapter contaminations. The average number of unique reads per library was 36931139, there were no low-quality base calls at reads' extremities with a peak of the distribution of the per read Phred quality score at 39. GC content was as expected with a peak at 51%, adapter content was less than 0.8% for all libraries.

Reads were mapped to the GRCh38 using the STAR aligner with ENCODE options:

--outFilterScoreMinOverLread 0.3 --outFilterMatchNminOverLread 0.3 --outFilterMultimapNmax 20 --alignSJoverhangMin 8 --alignSJDBoverhangMin 1 --outFilterMismatchNmax 999 --outFilterMismatchNoverReadLmax 0.04 --alignIntronMin 20 --alignIntronMax 1000000 --alignMatesGapMax 1000000

The median of uniquely aligned reads across libraries was 96.4 (IQR 96.2 - 96.6).

Gene counts were calculated using HTSeq against the the Grch38.88 gene set with the "intersection-nonempty" strategy. About 3% of reads (N = 1171455) were discarded as non-uniquely aligned during counting.

Differential Expression analysis was carried out in R using DESeq2. A model ~ Biological_Replicate + Well_Replicate + Group was fitted and DE was calculated using a bayesian zero-mean normal beta prior and the negative binomial wald test. Significance threshold was an FDR < 0.05.

Gene ontology (GO) analyses were performed in R using the GOseq package with a null distribution that considers count biases associated with the gene length.

HD10.6 were compared to human iPSC differentiated into sensory neurons from (Clark et al., 2021). Variance due to different library preparations was corrected as a batch effect using the limma "correctBatchEffect" function. The differentiation of iPSC to sensory neurons and UHD HD10.6 to neurons was preserved as an effect of interest.

Human neuronal subtype markers determined from Single-cell RNA-seq from human (Yu *et al.*, 2024) were used to draw heatmaps alongside neuronal marker genes from (Barry et al., 2023) . *PIEZO1* was substituted with *PIEZO2* as it is a more specific marker. Single cell human and mouse DRGs data from (Bhuiyan et al., 2024) was used to predict cell type proportions in bulk RNA-seq by doing deconvolution analysis with the SCDC R package. To do this HD10.6 Human RNA-seq data was mapped to mouse orthologs using biomaRt and the "findOrthologs" function (Vitalii Kleshchevnikov, 2020). Human and Mouse data was subsetted from the cross-species atlas and used separately.
