## Supplementary Figures and tables for "Extended maturation of the HD10.6 immortalised human dorsal root ganglion cell line enables modelling of nociceptive responses and neural injury"

### SUP FIG 1

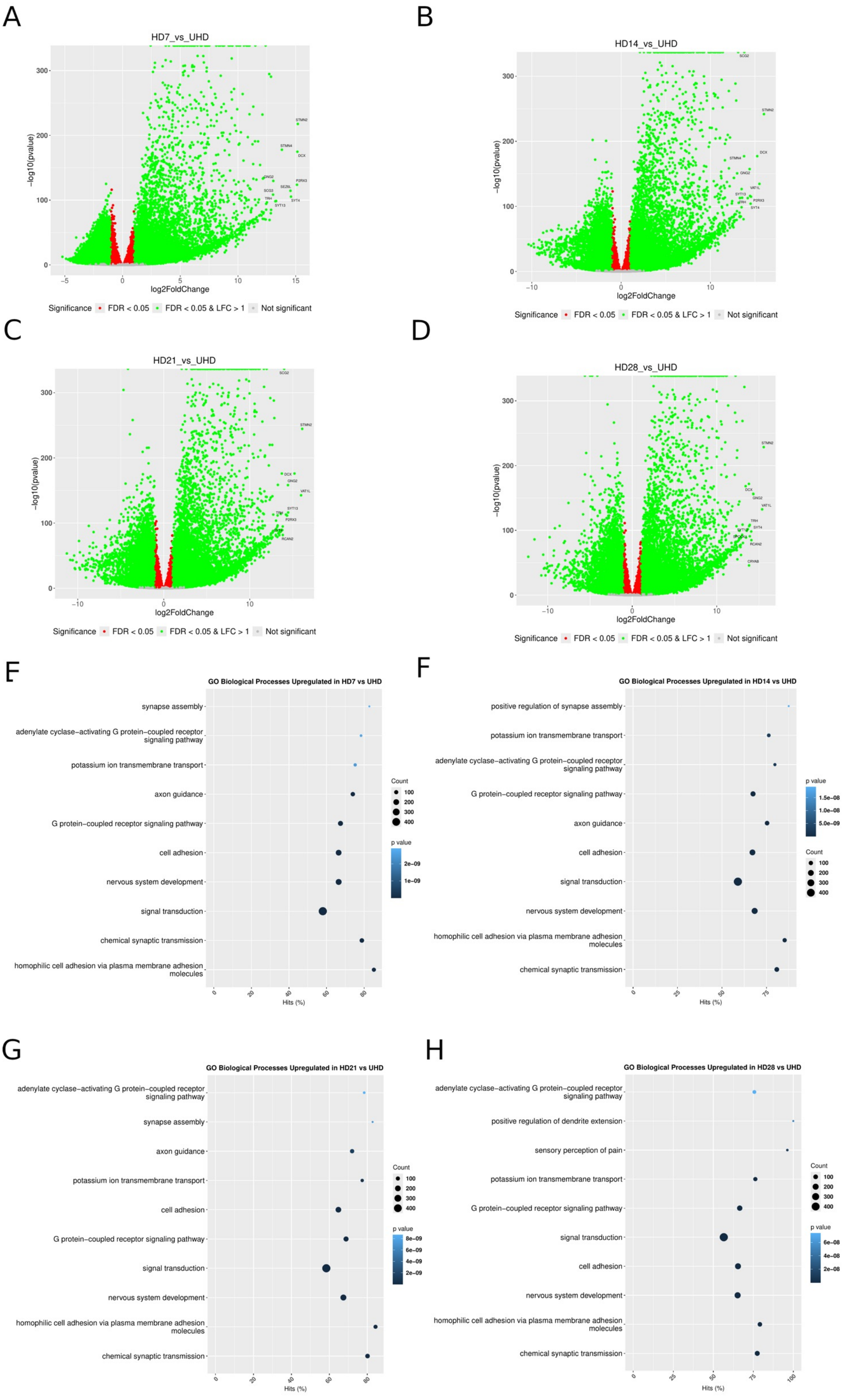
A
HD7_vs_UHD
-log10(pvalue)
log2FoldChange
STMN2
STMN4
DCX
GNG2
P2RX3
SCG3
SEZ6L
TRH
SYT4
SYT13
Significance FDR < 0.05 FDR < 0.05 & LFC > 1 Not significant
B
HD14_vs_UHD
-log10(pvalue)
log2FoldChange
SCG2
STMN2
DCX
STMN4
GNG2
VAT1L
SYT13
P2RX3
SYT4
Significance FDR < 0.05 FDR < 0.05 & LFC > 1 Not significant
C
HD21_vs_UHD
-log10(pvalue)
log2FoldChange
SCG2
STMN2
DCX
GNG2
VAT1L
SYT13
P2RX3
RCAN2
Significance FDR < 0.05 FDR < 0.05 & LFC > 1 Not significant
D
HD28_vs_UHD
-log10(pvalue)
log2FoldChange
STMN2
DCX
GNG2
VAT1L
TRH
SYT4
RCAN2
CRYAB
Significance FDR < 0.05 FDR < 0.05 & LFC > 1 Not significant
F
GO Biological Processes Upregulated in HD7 vs UHD
synapse assembly
adenylate cyclase-activating G protein-coupled receptor signaling pathway
potassium ion transmembrane transport
axon guidance
G protein-coupled receptor signaling pathway
cell adhesion
nervous system development
signal transduction
chemical synaptic transmission
homophilic cell adhesion via plasma membrane adhesion molecules
Hits (%)
Count
p value
F
GO Biological Processes Upregulated in HD14 vs UHD
positive regulation of synapse assembly
potassium ion transmembrane transport
adenylate cyclase-activating G protein-coupled receptor signaling pathway
G protein-coupled receptor signaling pathway
axon guidance
cell adhesion
signal transduction
nervous system development
homophilic cell adhesion via plasma membrane adhesion molecules
chemical synaptic transmission
Hits (%)
p value
Count
G
GO Biological Processes Upregulated in HD21 vs UHD
adenylate cyclase-activating G protein-coupled receptor signaling pathway
synapse assembly
axon guidance
potassium ion transmembrane transport
cell adhesion
G protein-coupled receptor signaling pathway
signal transduction
nervous system development
homophilic cell adhesion via plasma membrane adhesion molecules
chemical synaptic transmission
Hits (%)
Count
p value
H
GO Biological Processes Upregulated in HD28 vs UHD
adenylate cyclase-activating G protein-coupled receptor signaling pathway
positive regulation of dendrite extension
sensory perception of pain
potassium ion transmembrane transport
G protein-coupled receptor signaling pathway
signal transduction
cell adhesion
nervous system development
homophilic cell adhesion via plasma membrane adhesion molecules
chemical synaptic transmission
Hits (%)
Count
p value

### SUP FIG 2

A

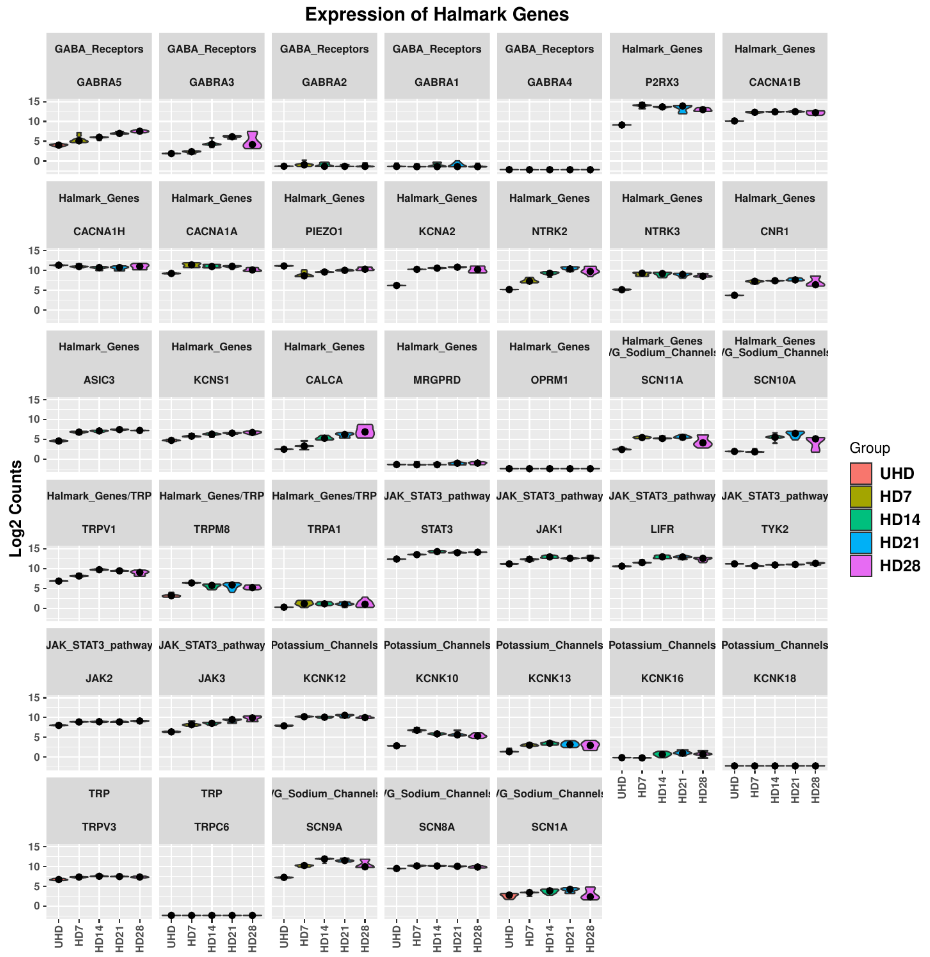

Expression of Halmark Genes
GABA_Receptors GABRA5
GABA_Receptors GABRA3
GABA_Receptors GABRA2
GABA_Receptors GABRA1
GABA_Receptors GABRA4
Halmark_Genes P2RX3
Halmark_Genes CACNA1B
Halmark_Genes CACNA1H
Halmark_Genes CACNA1A
Halmark_Genes PIEZO1
Halmark_Genes KCNA2
Halmark_Genes NTRK2
Halmark_Genes NTRK3
Halmark_Genes CNR1
Halmark_Genes ASIC3
Halmark_Genes KCNS1
Halmark_Genes CALCA
Halmark_Genes MRGPRD
Halmark_Genes OPRM1
Halmark_Genes /G_Sodium_Channel SCN11A
Halmark_Genes /G_Sodium_Channel SCN10A
Halmark_Genes/TRP TRPV1
Halmark_Genes/TRP TRPM8
Halmark_Genes/TRP TRPA1
JAK_STAT3_pathway STAT3
JAK_STAT3_pathway JAK1
JAK_STAT3_pathway LIFR
JAK_STAT3_pathway TYK2
JAK_STAT3_pathway JAK2
JAK_STAT3_pathway JAK3
Potassium_Channels KCNK12
Potassium_Channels KCNK10
Potassium_Channels KCNK13
Potassium_Channels KCNK16
Potassium_Channels KCNK18
TRP TRPV3
TRP TRPC6
/G_Sodium_Channel SCN9A
/G_Sodium_Channel SCN8A
/G_Sodium_Channel SCN1A
Log2 Counts
0 5 10 15
UHD HD7 HD14 HD21 HD28
Group
UHD
HD7
HD14
HD21
HD28


B

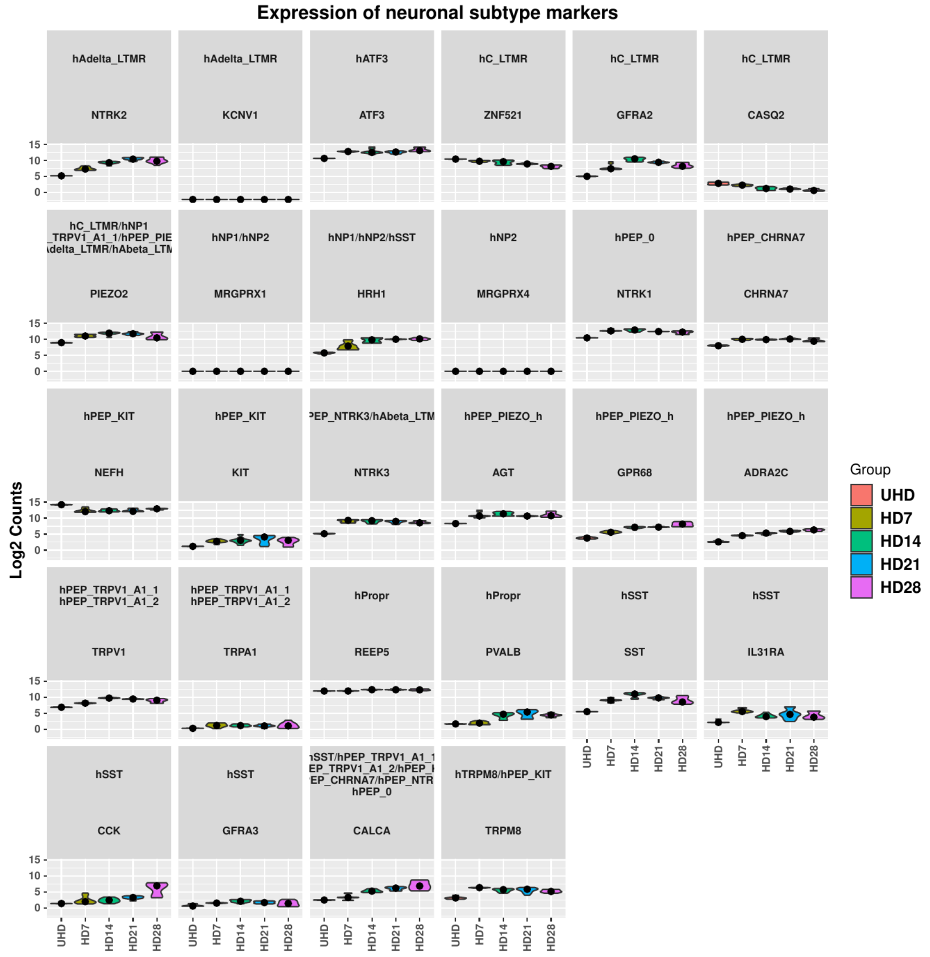

Expression of neuronal subtype markers
hAdelta_LTMR NTRK2
hAdelta_LTMR KCNV1
hATF3 ATF3
hC_LTMR ZNF521
hC_LTMR GFRA2
hC_LTMR CASQ2
hC_LTMR/hNP1 _TRPV1_A1_1/hPEP_PIE \delta_LTMR/hAbeta_LTM PIEZO2
hNP1/hNP2 MRGPRX1
hNP1/hNP2/hSST HRH1
hNP2 MRGPRX4
hPEP_0 NTRK1
hPEP_CHRNA7 CHRNA7
hPEP_KIT NEFH
hPEP_KIT KIT
'EP_NTRK3/hAbeta_LTM NTRK3
hPEP_PIEZO_h AGT
hPEP_PIEZO_h GPR68
hPEP_PIEZO_h ADRA2C
hPEP_TRPV1_A1_1 hPEP_TRPV1_A1_2 TRPV1
hPEP_TRPV1_A1_1 hPEP_TRPV1_A1_2 TRPA1
hPropr REEP5
hPropr PVALB
hSST SST
hSST IL31RA
hSST CCK
hSST GFRA3
\SST/hPEP_TRPV1_A1_1 EP_TRPV1_A1_2/hPEP_I EP_CHRNA7/hPEP_NTR hPEP_0 CALCA
hTRPM8/hPEP_KIT TRPM8
Log2 Counts
0 5 10 15
UHD HD7 HD14 HD21 HD28
Group
UHD
HD7
HD14
HD21
HD28

### SUP FIG 3

**A**

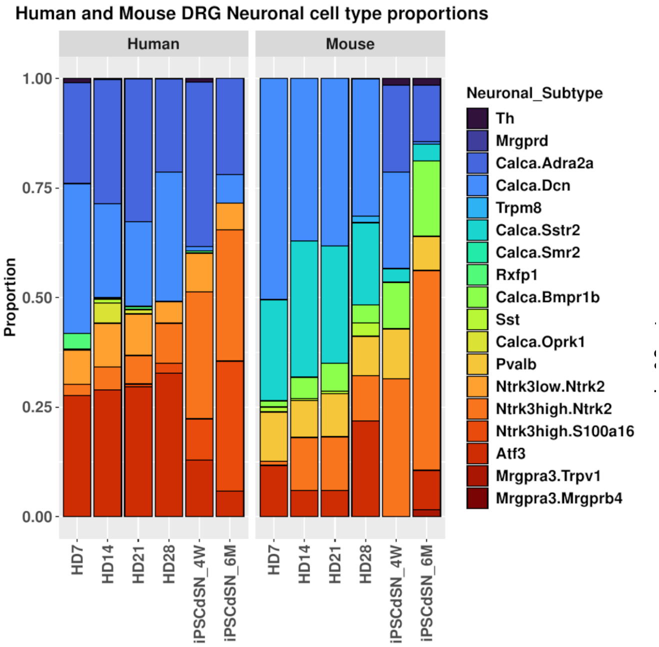
Human and Mouse DRG Neuronal cell type proportions
Human
Mouse
Proportion
1.00
0.75
0.50
0.25
0.00
HD7
HD14
HD21
HD28
iPSCdSN_4W
iPSCdSN_6M
Neuronal_Subtype
Th
Mrgprd
Calca.Adra2a
Calca.Dcn
Trpm8
Calca.Sstr2
Calca.Smr2
Rxfp1
Calca.Bmpr1b
Sst
Calca.Oprk1
Pvalb
Ntrk3low.Ntrk2
Ntrk3high.Ntrk2
Ntrk3high.S100a16
Atf3
Mrgpra3.Trpv1
Mrgpra3.Mrgprb4

**B**

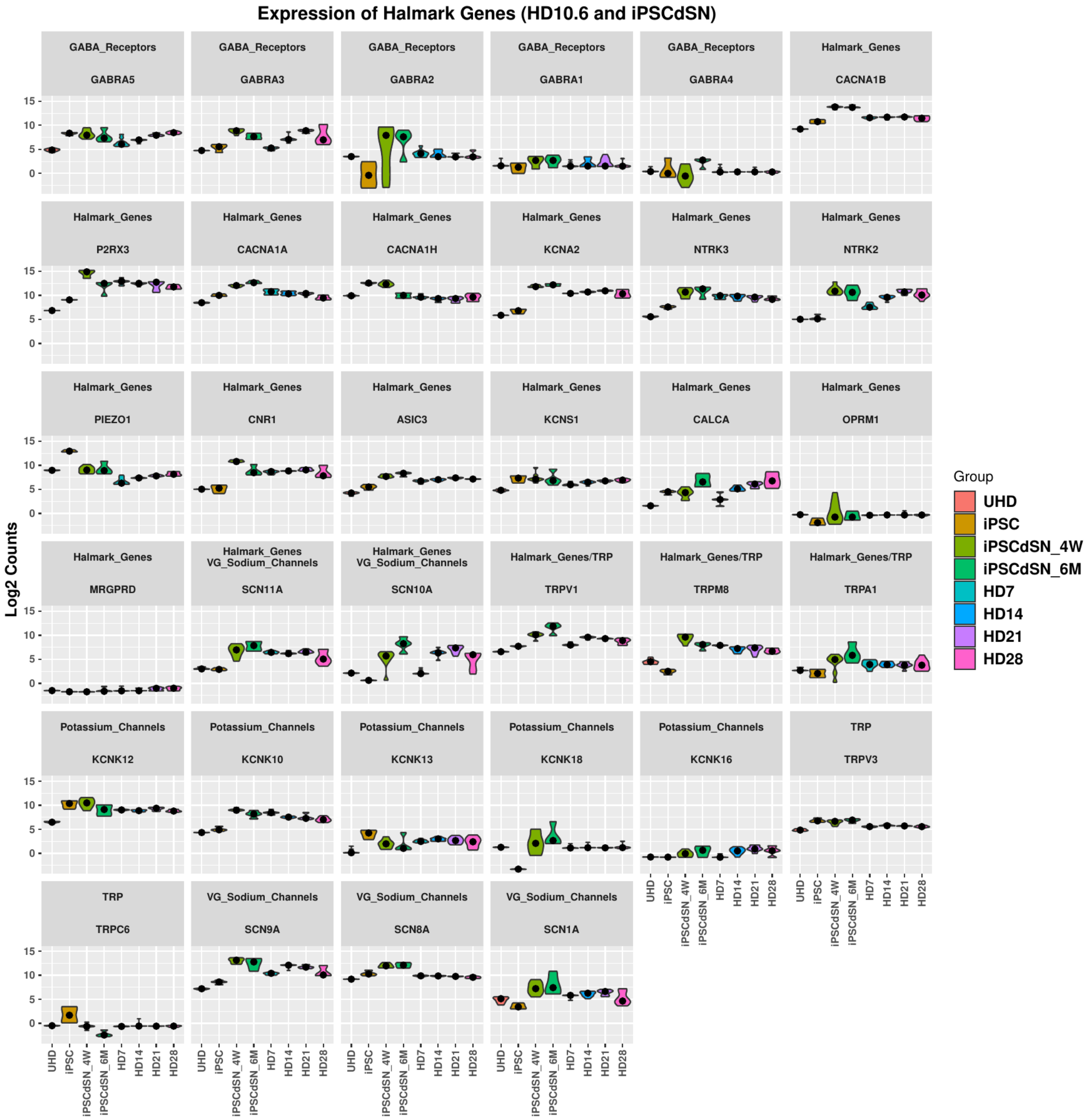
Expression of Halmark Genes (HD10.6 and iPSCdSN)
GABA_Receptors GABRA5
GABA_Receptors GABRA3
GABA_Receptors GABRA2
GABA_Receptors GABRA1
GABA_Receptors GABRA4
Halmark_Genes CACNA1B
Halmark_Genes P2RX3
Halmark_Genes CACNA1A
Halmark_Genes CACNA1H
Halmark_Genes KCNA2
Halmark_Genes NTRK3
Halmark_Genes NTRK2
Halmark_Genes PIEZO1
Halmark_Genes CNR1
Halmark_Genes ASIC3
Halmark_Genes KCNS1
Halmark_Genes CALCA
Halmark_Genes OPRM1
Halmark_Genes MRGPRD
Halmark_Genes VG_Sodium_Channels SCN11A
Halmark_Genes VG_Sodium_Channels SCN10A
Halmark_Genes/TRP TRPV1
Halmark_Genes/TRP TRPM8
Halmark_Genes/TRP TRPA1
Potassium_Channels KCNK12
Potassium_Channels KCNK10
Potassium_Channels KCNK13
Potassium_Channels KCNK18
Potassium_Channels KCNK16
TRP TRPV3
TRP TRPC6
VG_Sodium_Channels SCN9A
VG_Sodium_Channels SCN8A
VG_Sodium_Channels SCN1A
Log2 Counts
15
10
5
0
UHD
iPSC
iPSCdSN_4W
iPSCdSN_6M
HD7
HD14
HD21
HD28
Group
UHD
iPSC
iPSCdSN_4W
iPSCdSN_6M
HD7
HD14
HD21
HD28

### SUP FIG 4

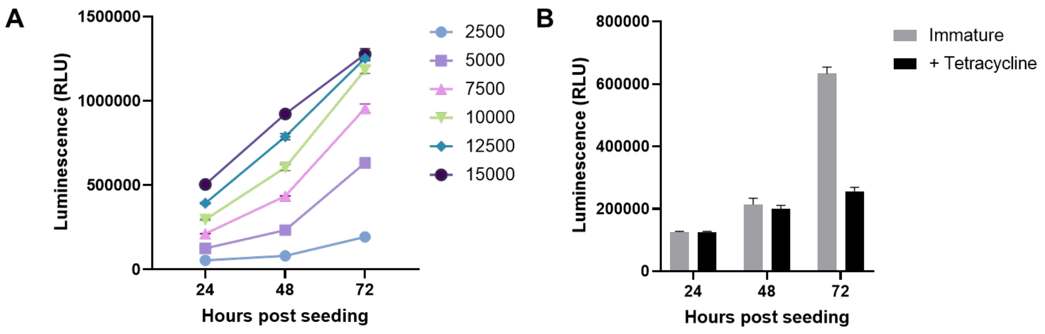
A
1500000
1000000
500000
0
Luminescence (RLU)
24
48
72
Hours post seeding
2500
5000
7500
10000
12500
15000
B
800000
600000
400000
200000
0
Luminescence (RLU)
24
48
72
Hours post seeding
Immature
+ Tetracycline

### SUP FIG 5

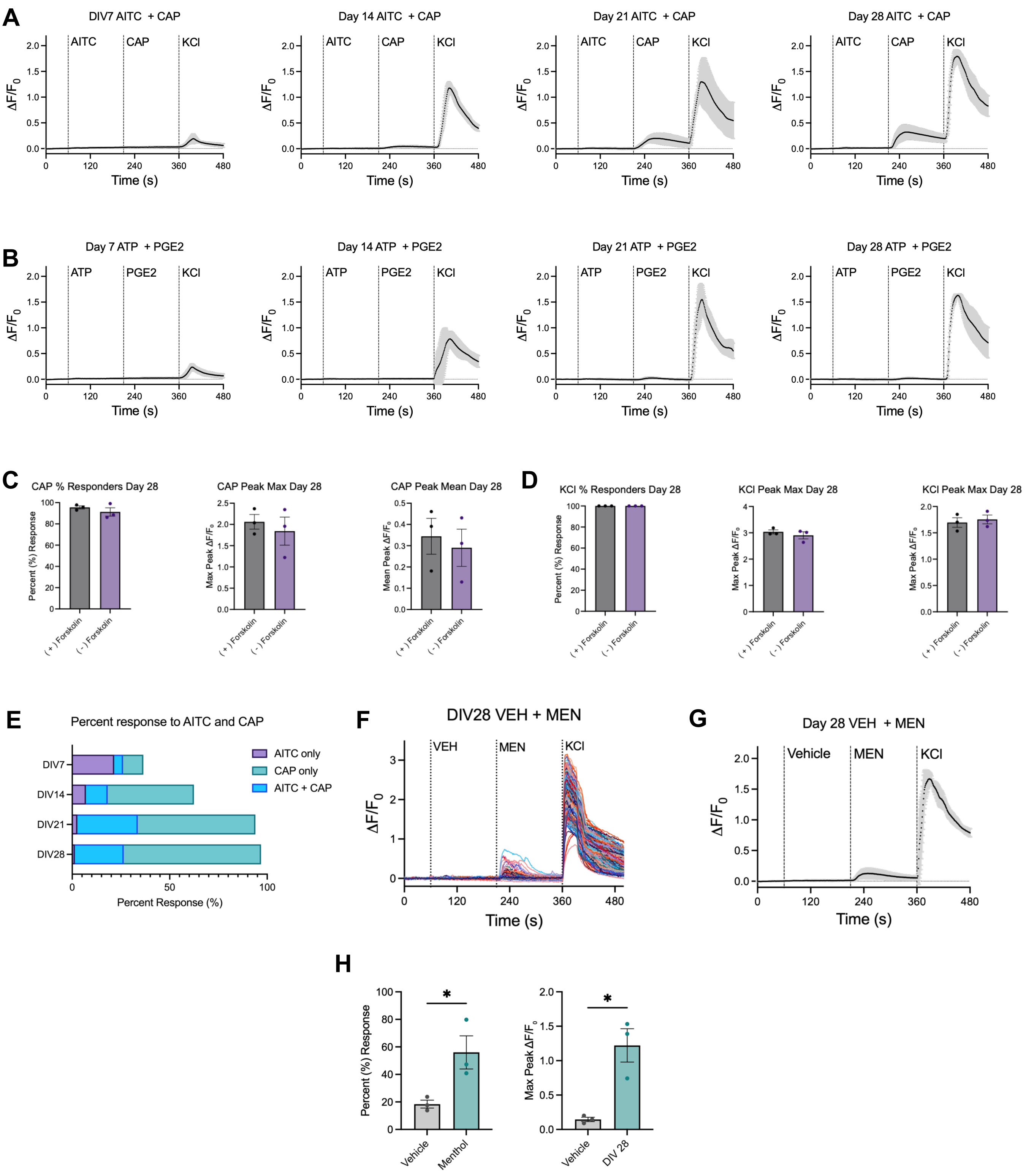
A
DIV7 AITC + CAP
Day 14 AITC + CAP
Day 21 AITC + CAP
Day 28 AITC + CAP
AITC
CAP
KCl
ΔF/F0
Time (s)
B
Day 7 ATP + PGE2
Day 14 ATP + PGE2
Day 21 ATP + PGE2
Day 28 ATP + PGE2
ATP
PGE2
C
CAP % Responders Day 28
CAP Peak Max Day 28
CAP Peak Mean Day 28
Percent (%) Response
Max Peak ΔF/F0
Mean Peak ΔF/F0
(+) Forskolin
(-) Forskolin
D
KCl % Responders Day 28
KCl Peak Max Day 28
KCl Peak Max Day 28
E
Percent response to AITC and CAP
DIV7
DIV14
DIV21
DIV28
AITC only
CAP only
AITC + CAP
Percent Response (%)
F
DIV28 VEH + MEN
VEH
MEN
G
Day 28 VEH + MEN
Vehicle
H
Vehicle
Menthol
DIV 28
*

SUP FIG 6

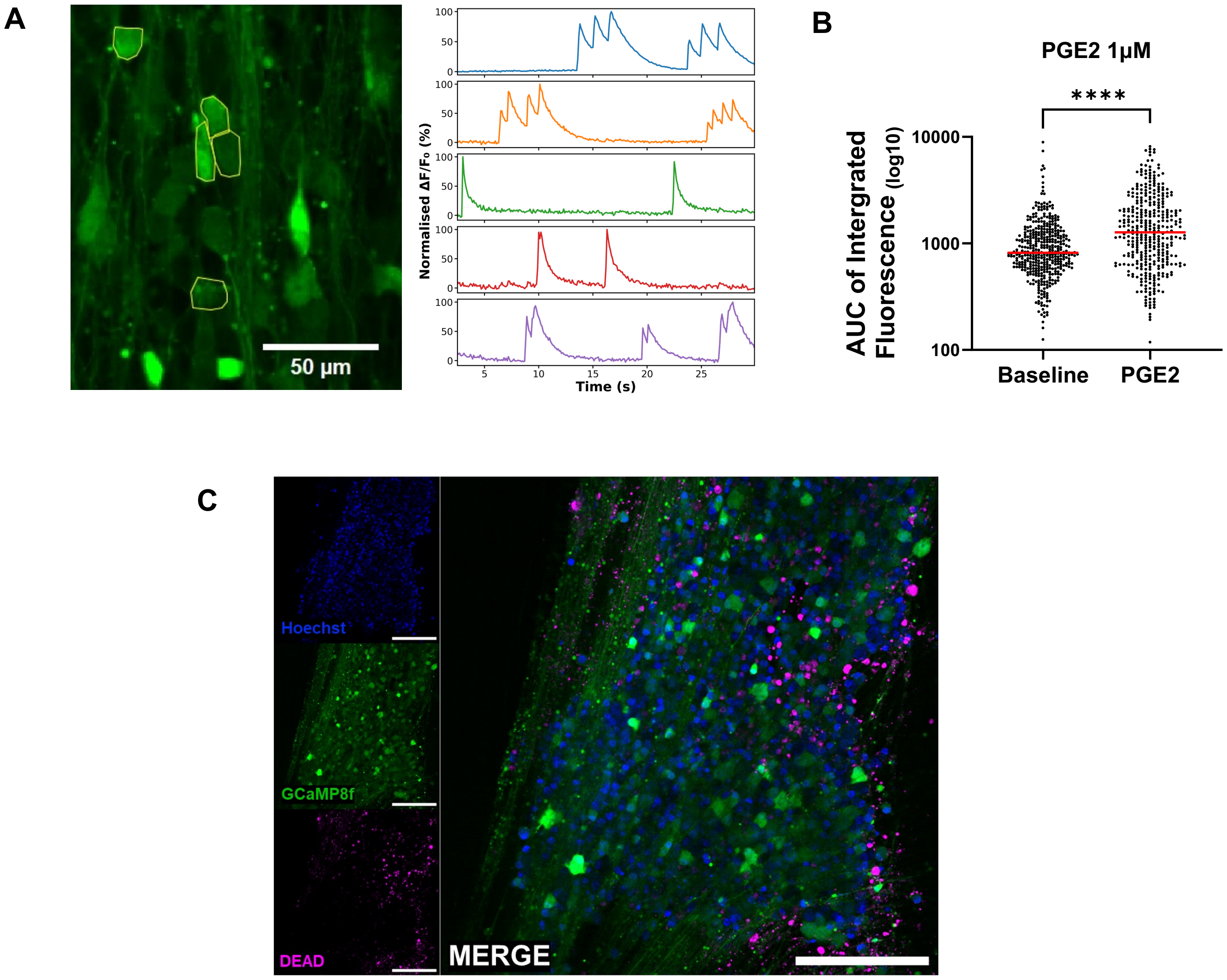
A
50 µm
Normalised ΔF/F₀ (%)
Time (s)
B
PGE2 1µM
****
AUC of Intergrated Fluorescence (log10)
10000
1000
100
Baseline
PGE2
C
Hoechst
GCaMP8f
DEAD
MERGE

SUP FIG 7

A

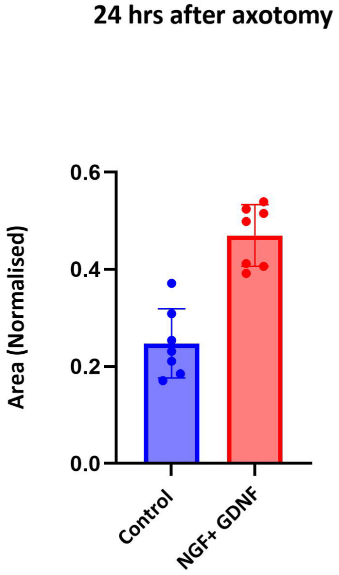
24 hrs after axotomy
Area (Normalised)
0.6
0.4
0.2
0.0
Control
NGF+ GDNF

SUP FIG 8

**A**

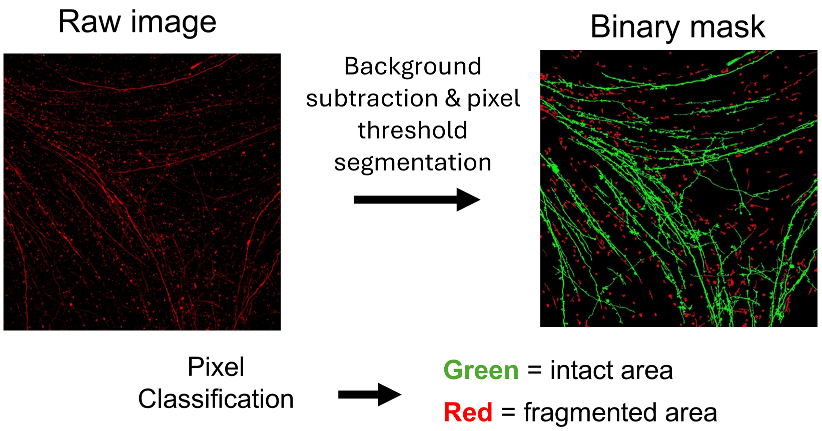


**B**

$$Fragmentation\ index = \frac{fragmented\ area}{fragmented\ area + intact\ area}$$

### SUP Table 1

#### Table S1. Antibodies and concentrations

| **Target** | **Host / type** | **Use** | **Concentration** | **Manufacturer / Cat.** |
|---|---|---|---|---|
| NF200 | Mouse monoclonal | ICC | 1:200 | Sigma (N0142) |
| BIII-Tubulin | Rabbit monoclonal | ICC | 1:1000 | Abcam (ab202519) |
| Peripherin | Mouse monoclonal | ICC | 1:250 | Santa Cruz (sc-377093) |
| CGRP | Sheep polyclonal | ICC | 1:250 | Enzo (BML-CA1137) |
| BRN3a | Rabbit polyclonal | ICC | 1:500 | Sigma (AB5945) |
| BRN3a | Rabbit polyclonal | WB | 1:200 | Bioss (BS-3669R) |
| NaV 1.7 | Mouse monoclonal | WB | 1:500 | Abcam (ab85015) |
| NaV 1.8 | Rabbit polyclonal | ICC / WB | 1:200 / 1:200 | Alomone (AS-016) |
| Beta- Actin | Mouse monoclonal | WB | 1:5000 | Invitrogen (MA5-15739) |
| TRPV1 | Rabbit polyclonal | ICC / WB | 1:200 / 1:200 | Sigma (SAB5700857) |
| Krox-20 | Rabbit polyclonal | ICC | 1:300 | Convance (PRB-236P-100) |
| Myelin Basic Protein | Rat monoclonal | ICC | 1:300 | Abcam (ab7349) |
| SARM1 | Rabbit monoclonal | WB | 1:500 | Abcam (ab309195) |
| DAPI | - | ICC | 1 µg / mL | Invitrogen (D13) |

### SUP Table 2

| Name | Final concentration | Manufacturer and cat. |
|---|---|---|
| Fibronectin | 16.6 µg/ml final | Milipore Sigma, FC010 |
| Laminin | 1:300 | Milipore Sigma, L2020 |
| Poly-D-Lysine | 1:100 | Milipore Sigma, P6407 |
| Cultrex | 1:100 | R&D systems, 3434-005-02 |
| Advanced DMEM/F12 | Base medium | Gibco, 12634-010 |
| B27 Supplement | 1× | Gibco, 17504-044 |
| GlutaMAX | 1× | Gibco, 35050061 |
| Prostaglandin E1 | 10 ng/ml | Sigma, P5515 |
| bFGF | 0.5 ng/ml | PeproTech, 100-18B |
| G418 Solution | 50 µg/ml | Roche, 04727878001 |
| Neurobasal™ Medium, minus phenol red | Base medium | Gibco, 12348017 |
| NGF | 50 ng/ml | R&D Systems, 256-GF |
| CNTF | 25 ng/ml | PeproTech, #450-13 |
| GDNF | 25 ng/ml | R&D Systems, 217-GD |
| NT-3 | 25 ng/ml | R&D Systems, 267-N3 |
| Tetracycline | 1 µg/ml | Sigma, T-7660 |
| Forskolin | 25 µM | Sigma, F6886 |

#### Supplementary Figure legends

##### Supplementary Figure 1: RNA-seq analysis identifies progressive acquisition of neuronal and sensory biological programmes during HD10.6 maturation

**(A–D)** Volcano plots showing differential gene expression in HD10.6 sensory neurons differentiated for 7 days (HD7), 14 days (HD14), 21 days (HD21) and 28 days (HD28) relative to undifferentiated HD10.6 cells (UHD). The x-axis shows $\log_2$ fold change and the y-axis shows $-\log_{10}$ (adjusted P-value). Differentially expressed genes passing the indicated thresholds are highlighted, with significantly upregulated and downregulated transcripts distinguished from non-significant genes.
**(E–H)** Gene Ontology (GO) biological process enrichment analysis of genes upregulated in HD7, HD14, HD21 and HD28, respectively, relative to UHD. Dot size indicates gene count, colour indicates P-value, and the x-axis shows hits (%).

##### Supplementary Figure 2: Temporal expression of hallmark sensory neuronal genes and subtype-associated markers during HD10.6 maturation

**(A)** Expression of selected hallmark genes across immature HD10.6 cells (UHD) and HD10.6 sensory neurons differentiated for 7, 14, 21 and 28 days (HD7, HD14, HD21 and HD28).
**(B)** Expression of individual values of selected neuronal subtype markers across UHD, HD7, HD14, HD21 and HD28, plotted as $\log_2$ counts. Markers shown represent transcriptionally defined sensory neuronal subtypes (determined in Yu et al., 2024).

##### Supplementary Figure 3: Cross-species deconvolution and hallmark gene expression profiling of HD10.6 sensory neurons and iPSCdSNs

**(A)** Predicted proportions of separated human and mouse dorsal root ganglion (DRG) neuronal subtypes (determined by (Bhuiyan et al., 2024) across immature HD10.6 cells (UHD), HD10.6 sensory neurons matured for 7, 14, 21 and 28 days (HD7, HD14, HD21 and HD28), and human iPSC-derived sensory neurons matured for 4 weeks (iPSCdSN_4W) or 6 months (iPSCdSN_6M). Stacked bar plots show the relative contribution of transcriptionally defined neuronal subtypes, including populations associated with nociceptor, mechanoreceptor and proprioceptor identities.
**(B)** Expression of selected hallmark genes across UHD, iPSCdSN_4W, iPSCdSN_6M, HD7, HD14, HD21 and HD28.

**Supplementary Figure 4: Cell Titre Glo ATP viability measurements to show and quantify proliferation**

**(A)** Luminescence (RLU) of increasing density (cells per well) of immature HD10.6 cells at 24, 48 and 72 hours after seeding (n = 5-6, N = 1)

**(B)** Luminescence (RLU) of immature HD10.6 cells (5000 cells per well) at 24, 48 and 72 hours after seeding with and without treatment with 1 μM tetracycline (media exchanged at 24 hours after seeding).

**Supplementary Figure 5: Agonist-evoked calcium responses in DIV28 HD10.6 sensory neurons**

**(A)** Mean calcium responses ($\Delta F/F_0$) of matured HD10.6 neurons at DIV7, DIV14, DIV21 and DIV28 following sequential application of 50 μM allyl isothiocyanate (AITC), 1 μM capsaicin (CAP) and 50 mM KCl, with timings indicated by vertical dashed lines.

**(B)** Representative calcium responses at DIV7, DIV14, DIV21 and DIV28 following sequential application of 100 μM α,β-MeATP, 1 μM prostaglandin E2 (PGE2) and 50 mM KCl.

**(C)** Percentage of responding cells, maximal peak and mean peak $\Delta F/F_0$ response to capsaicin (1 μM) in DIV28 HD10.6 sensory neurons maintained with (+) or without (−) forskolin. Data shows ± SEM. Wilcoxon (non-parametric) test used to compare paired means of groups from 3 separate biological replicates (N = 3)

**(D)** Percentage of responding cells, maximal peak and mean peak $\Delta F/F_0$ response to to KCl (50 mM) in DIV28 HD10.6 sensory neurons maintained with (+) or without (−) forskolin.

**(E)** Proportion of AITC- and capsaicin-responsive neuronal populations at DIV7, DIV14, DIV21 and DIV28, separated into AITC-only (purple), capsaicin-only (teal) and dual AITC+CAP (blue) responders.

**(F)** Representative calcium traces from DIV28 HD10.6 sensory neurons following sequential application of ethanol vehicle, 100 μM menthol (MEN) and 50 mM KCl.

**(G)** Mean calcium traces from DIV28 HD10.6 sensory neurons following sequential application of ethanol vehicle, 100 μM menthol (MEN) and 50 mM KCl,. (n = 1480 cells, from 3 independent maturations).

**(H)** Quantification of menthol-evoked responses at DIV28, shown as percentage of responding cells and maximal peak $\Delta F/F_0$ compared to ethanol vehicle. Data shows ± SEM.

Mean traces show mean ± SD. Responding cells threshold: > 3 x SD of pre-30 seconds baseline. For quantification graphs, bars show mean ± SEM and dots represent individual biological replicates/independent experiments. Statistical comparisons are indicated on the graphs; *$P < 0.05$, **$P < 0.01$, ***$P < 0.001$, ****$P < 0.0001$.

**Supplementary Figure 6: GCaMP8f calcium imaging after treatment with Prostaglandin E2 and LIVE/DEAD validation.**

**(A)** Representative calcium imaging field from differentiated neurons with example ROIs from cells (left) and corresponding single-cell normalised (%) $\Delta F/F_0$ traces (right), where $F_0$ is 8% percentile of baseline, illustrating spontaneous calcium transients. Scale bar = 50 μm.

**(B)** Single-cell analysis showing integrated fluorescence (area under curve of baseline-subtracted $\Delta F/F_0$) over 30 s (10 Hz) of DIV32 HD10.6 neuronal cells imaged at baseline, and right after treatment with PGE2 (1 μM). Each point represents one cell; red line shows median. Kruskal–Wallis test (nonparametric one-way ANOVA; $p < 0.0001$; N = 3).

**(C)** Representative live image showing GCaMP8f positive cells (green), dead cells (magenta) and Hoechst (blue) nuclear marker of HD10.6 sensory neurons

**Supplementary Figure 7: Quantification of Regeneration**

**(A)** Quantification of neurite regeneration expressed as the area occupied by EYFP-positive neurites, normalised to pre-axotomy values. NGF/GDNF treatment increased neurite regeneration compared to control conditions lacking growth factors.

**Supplementary Figure 8: Fragmentation index**

**(A)** Representative images of segmentation of raw live image of TdTomato positive HD10.6 neurons, and after pixel size thresholding to determine fragmented (red) and unfragmented (green) neurites for calculation of fragmentation index.

**(B)** Equation used to calculate Fragmentation Index
